# Optogenetic restoration of neuron subtype-specific cortical activity ameliorates motor deficits in Huntington’s Disease mice

**DOI:** 10.1101/2025.02.07.637155

**Authors:** Sonja Blumenstock, David Arakelyan, Nicholas del Grosso, Sonja Schneider, Yufeng Shao, Enida Gjoni, Rüdiger Klein, Irina Dudanova, Takaki Komiyama

**Affiliations:** Department of Neurobiology, Center for Neural Circuits and Behavior, and Department of Neurosciences, University of California San Diego, La Jolla, CA 92093, USA; Department of Molecules – Signaling – Development, Max Planck Institute for Biological Intelligence, 82152 Martinsried, Germany; Molecular Neurodegeneration Group, Max Planck Institute for Biological Intelligence, 82152 Martinsried, Germany; Center for Anatomy, Faculty of Medicine and University Hospital Cologne, University of Cologne, 50931 Cologne, Germany; Cologne Excellence Cluster on Cellular Stress Responses in Aging-Associated Diseases (CECAD), University of Cologne, 50931 Cologne, Germany; Halıcıoğlu Data Science Institute, University of California San Diego, La Jolla, CA 92093, USA

## Abstract

Huntington’s disease (HD) is a devastating movement disorder without a current cure. Although the monogenic basis of HD is well-defined, the complex downstream effects that underlie behavioral symptoms are poorly understood. These effects include cortical dysfunctions, yet the role of specific cortical neuronal subtypes in HD symptoms remain largely unexplored. Here, we used longitudinal *in vivo* two-photon calcium imaging to examine the activity of two cortical inhibitory neuron (IN) subtypes and excitatory corticostriatal projection neurons (CSPNs) in the motor cortex of R6/2 HD mouse model throughout disease progression. We found that motor deficits in R6/2 mice were accompanied by neuron type-specific abnormalities in movement-related activity, including hypoactivity of vasoactive intestinal peptide (VIP)-INs and CSPNs. Optogenetic activation of VIP-INs in R6/2 mice restored healthy levels of activity in VIP-INs and their downstream CSPNs and ameliorated motor deficits in R6/2 mice. Our findings highlight cortical INs as a potential therapeutic target for HD and possibly other neurological diseases.

## Introduction

Precise motor control is essential for an animal’s survival and well-being. This function is subserved by complex, interconnected neuronal networks in the brain that include the neocortex and striatum^1^. Various neurodegenerative diseases affect motor control, including Huntington’s disease (HD). HD is a devastating monogenic disorder without a current cure, caused by a CAG repeat expansion in the Huntingtin (HTT) gene^2^. The disease is characterized by a severe and progressive motor dysfunction. While the genetic foundation of HD is well understood^2^, the precise neural circuit basis that leads to HD motor symptoms remains unclear. Notably, one of the key brain regions associated with the symptomatology and progression of HD is the neocortex and its projections to the striatum. Changes in corticostriatal connections are among the earliest events in HD progression, and disconnection from cortical afferents likely plays a major role in the subsequent dysfunction of the downstream striatal circuits^3–5^. Furthermore, cortical inhibitory circuits, which regulate the activity of excitatory corticostriatal projection neurons (CSPNs), appear to be impacted in HD^6–8^. These observations raise the possibility that HD has cell type-specific effects on cortical networks. Many neuron types are molecularly identifiable and genetically targetable, with the potential for subtype-specific therapeutic interventions.

Cortical inhibitory neurons (INs) consist of distinct subtypes with unique functions shaped by their morphology, connectivity, and physiology. Among these subtypes, those expressing vasoactive intestinal peptide (VIP) and somatostatin (SST) play important roles in regulating cortical circuits, with SST-INs mainly inhibiting excitatory neuron dendrites, and VIP-INs mainly inhibiting other INs^9–12^. The precise regulation of the activity of these functionally diverse subtypes is essential for proper cortical circuit functions underlying healthy behavior. However, the impact of HD on distinct cortical IN subtypes has been poorly explored, and direct evidence of the pathogenic interplay between cortical inhibition and CSPNs is lacking.

Here we used longitudinal *in vivo* two-photon calcium imaging to examine the activity of VIP- and SST-INs as well as CSPNs in the motor cortex in HD model mice during spontaneous and learned movements throughout the disease progression. We uncovered a neuron subtype-specific dysfunction of cortical circuits in HD. We further demonstrate that neuron subtype-specific activity modulation using optogenetics can normalize cortical network activity and ameliorate motor deficits in HD mice.

## Results

### Motor learning deficits in HD mice are reflected by subtype-specific alteration of cortical neuron activity

To investigate the neural mechanisms underlying motor symptoms in HD, we used the transgenic R6/2 mouse model of HD, which expresses exon 1 of the human mutant huntingtin (*mHTT*) gene and is characterized by motor impairments and reduced life span^13^. Non-transgenic littermates were used as controls throughout. We trained mice on a motorized ladder task, in which mice learned to walk on a circular ladder with equally spaced rungs under head-fixation in the dark. Following an auditory cue, the ladder rotated at a fixed speed for 8 seconds in each trial, during which mice had to walk on the ladder by grasping the rungs. After a 5-day training regimen, mice were repeatedly tested in the task, covering early, middle, and late stages in learning / disease progression (**Fig. 1a-b, Extended Data Fig. 1a, Methods**). During the task, body movements were traced by video analysis^14^ (**Extended Data Fig. 1b**). Compared to controls, R6/2 mice exhibited a range of deficits in movement kinematics (**Fig. 1c**), and we analyzed a number of kinematic features to characterize the deficits. First, we quantified hindlimb dragging as a measure of failures to keep up with the rotation of the ladder (**Extended Data Fig. 1c**). R6/2 mice exhibited more hindlimb dragging compared to ctrl mice, with the difference becoming more apparent over time (**Fig. 1d**). Next, we assessed the regularity of the gait pattern by an autocorrelation analysis of paw trajectories, which revealed a consistently lower regularity in the gait pattern of R6/2 mice (**Fig. 1e, Extended Data Fig. 1d**). We then analyzed the distribution of cadence during gait, using Fourier analysis of paw trajectories. While control mice showed a clear peak of cadence at ~1.3 Hz, indicating consistency of stride frequency, R6/2 mice lacked such a peak and showed overall reduced cadence (**Fig. 1f, Extended Data Fig. 1e**). In addition, the directness of strides—defined as the linearity of the forepaw trajectories during the swing phase of locomotion — was reduced in R6/2 mice and further deviated from ctrl mice with disease progression (**Fig. 1g**). These various measures confirm motor deficits in R6/2 mice.

**Fig. 1:**
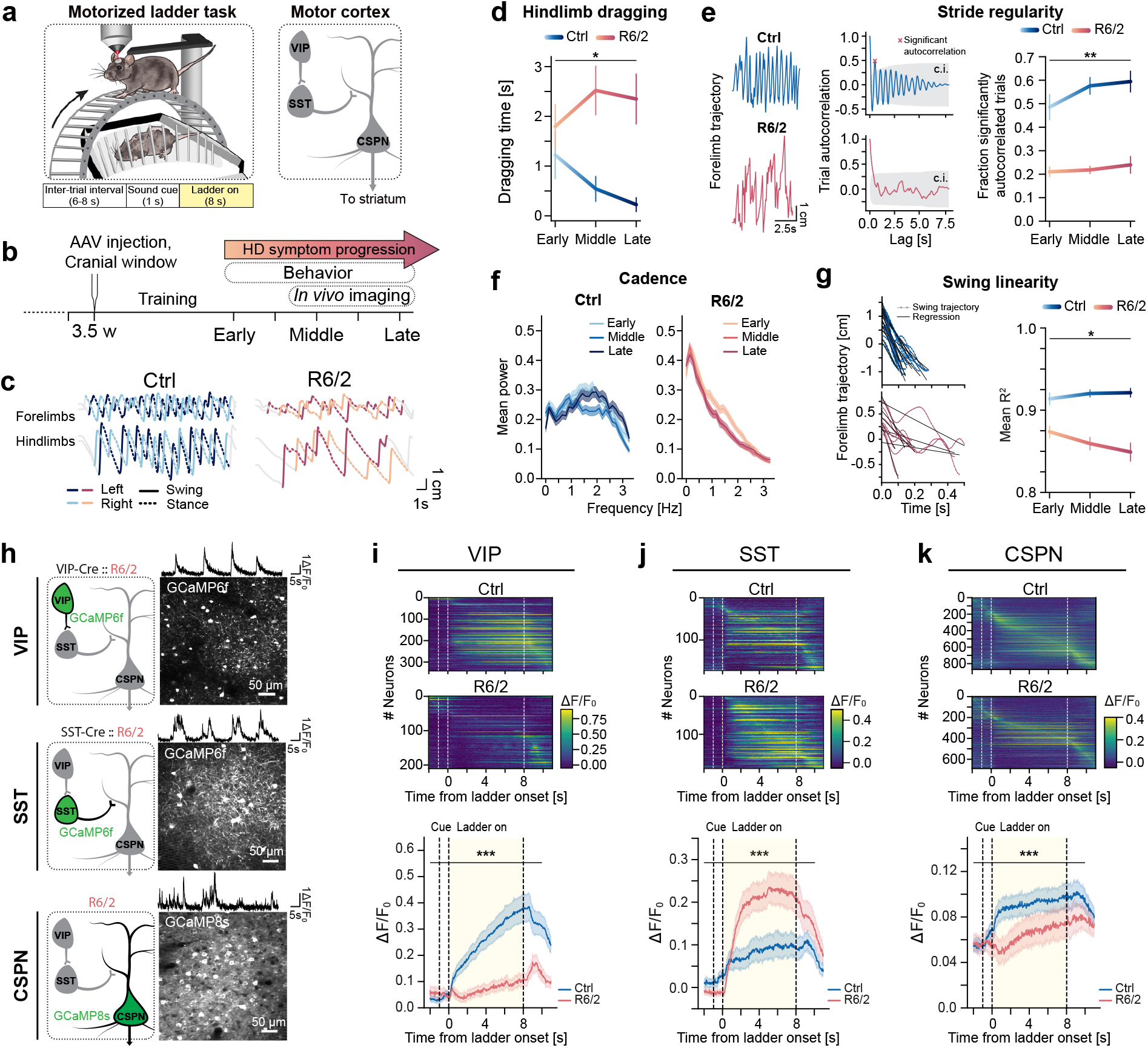
Motor dysfunction in HD mice is reflected by subtype-specific alteration of cortical neuron activity. (**a**) Left, schematic of the experimental setup and task structure. Right, simplified connectivity scheme of the examined neuron subtypes. (**b**) Diagram of behavioral and imaging schedule. Early, middle and late timepoints were defined through body weight and CAG repeat numbers of the R6/2 mice (Extended Data Fig. 1a, Methods). (**c**) Example movement trajectories of forelimbs and hindlimbs in single trials in ctrl and R6/2 mice at middle timepoint sessions. (**d**) Duration of hindlimb dragging per trial was longer in R6/2 mice and progressively deviated from ctrls with advancing timepoints (Aligned rank transform; genotype main effect, *p* = 0.012; genotype:timepoint interaction, *p* = 0.017). (**e**) Regularity of strides measured by the autocorrelation of forelimb trajectories. Left, example trajectories in single trials at the middle timepoint. Middle, autocorrelation plots from example trials. Right, fractions of significantly autocorrelated trials. R6/2 mice showed lower fractions of significantly autocorrelated trials, indicating their irregularity of gait pattern (Aligned rank transform; genotype main effect, *p* < 0.001; genotype:timepoint interaction, *p* = 0.005). c.i., 95% confidence interval. (**f**) Cadence of strides was measured by Fourier analysis. Average power spectra of trial forelimb trajectories. Cadence in ctrl mice was consistent around 1-2 steps per second. R6/2 mice step frequency was inconsistent. (**g**) The linearity of strides was measured by linear regression of the swing phase of single forelimb strides. Left: strides of single example trials, aligned to the swing onset of the stride phase, with overlaid regression lines. Right: mean R^2^ values, representing a measure for the directness of swings. R6/2 swings were less direct than ctrl swings and further deviated from ctrls over timepoints (Aligned rank transform; genotype main effect, *p* < 0.001; genotype:timepoint interaction, *p* = 0.037). (d-g) n = 8 ctrl, n = 8 R6/2 mice, n = 107 sessions across all mice and timepoints. (**h**) Example maximum intensity projections of *in vivo* two-photon fluorescence images and example calcium traces from individual neurons. VIP- and SST-INs expressed Cre-dependent GCaMP6f (in VIP-Cre::R6/2 and SST-Cre::R6/2 mice, respectively), and CSPNs were labeled retrogradely with GCaMP8s from the dorsolateral striatum. Imaging was performed at middle and late timepoints. **(i)** Calcium signals of VIP-INs (n = 342 neurons / 17 sessions from 8 VIP-Cre ctrl mice, 211 neurons / 16 sessions from 8 VIP-Cre::R6/2 mice) averaged over trials, from 2 s before ladder onset to 3 s after ladder offset. Data was binned into 2 s intervals for statistical comparison (Aligned rank transform; genotype main effect, *p* = 0.018; genotype:time bin interaction, *p* < 0.001). **(j)** Calcium signals from SST-INs (n = 173 neurons / 15 sessions from 4 SST-Cre ctrl mice, n = 191 neurons / 19 sessions from 4 SST-Cre::R6/2 mice; Aligned rank transform; genotype main effect, *p* = 0.008; genotype:time bin interaction, *p* < 0.001). **(k)** Calcium signals from CSPNs (n = 871 neurons / 18 sessions from 6 ctrl mice, n = 689 neurons / 18 sessions from 5 R6/2 mice; Aligned rank transform; genotype main effect, *p* = 0.001; genotype:time bin interaction, *p* < 0.001). For this and all other figures, *: p<0.05; **: p<0.01; ***: p<0.001 for interaction effects, and #: p<0.05; ##: p<0.01; ###: p<0.001 for group effects. Error bars represent mean ± s.e.m for behavioral analyses and mean ± c.i. for neuronal activity measurements.

To explore potential cortical contributions underlying these behavioral deficits, we recorded the activity of several cortical neuron subtypes in mice during the ladder task with two-photon calcium imaging. We labeled VIP- and SST-INs by injecting AAV vectors expressing GCaMP6f in a Cre-dependent manner into the primary motor cortex (M1) of VIP-Cre and SST-Cre mice^15^, crossed to R6/2 mice, respectively (**Fig. 1b, h**). In R6/2 mice, VIP-INs exhibited a nearly complete absence of movement-related activity (**Fig. 1i**). In contrast, SST-INs were hyperactive during movements in R6/2 mice compared to the controls (**Fig. 1j**). To investigate how such subtype-specific dysregulations of cortical INs in R6/2 mice may be reflected in the activity of cortical principal neurons, we next examined CSPNs projecting to the dorsolateral striatum, which is the part of striatum most severely affected in HD. These CSPNs were targeted by retrograde labeling to express GCaMP8s. By imaging CSPNs during the ladder task, we observed that their activity during movement was significantly reduced in R6/2 mice compared to the controls (**Fig. 1k**). These results reveal neuron subtype-specific dysfunctions of the cortical circuit in HD. In particular, considering the known connectivity among these three neuron subtypes (VIP-INs inhibit SST-INs which in turn inhibit CSPNs, **Fig. 1a**), the absence of movement-related activity in VIP-INs could be partially responsible for the hyperactivity and hypoactivity of SST-INs and CSPNs, respectively.

### Abnormalities of spontaneous movements in HD mice are accompanied by subtype-specific dysfunctions of cortical INs

To test whether the altered activity of cortical IN subtypes is generalizable across different movements, we employed another behavioral paradigm where head-fixed mice were placed on a continuous-surface wheel that mice could turn by spontaneous locomotion. This was done repeatedly for early, middle, and late phases as described above (**Extended Data Fig. 2a, b**). We did not impose any task structure and instead observed the spontaneous behavior of the mice examined by video analysis (**Fig. 2a**). Using a semi-supervised deep learning algorithm^16^ applied on videos, behavioral epochs were classified into different behavioral categories (**Fig. 2a, Extended Data Fig. 2c**). These categories of spontaneous behaviors included ‘active’ epochs such as ‘locomotion’, ‘grooming’, and forelimb ‘twitches’, as well as ‘inactive’ epochs corresponding to periods labeled as ‘rest’ and ‘sit’ (Methods). Control mice showed a distribution of these epochs that were consistent throughout the timepoints, while R6/2 mice demonstrated a different distribution at the early stage as well as a progressive deviation from the pattern seen in the control mice. Specifically, the duration of active epochs significantly decreased and inactive epochs correspondingly increased (**Fig. 2b**). The decrease of active epochs in R6/2 mice was largely accounted for by a decrease in locomotion periods.

**Fig. 2:**
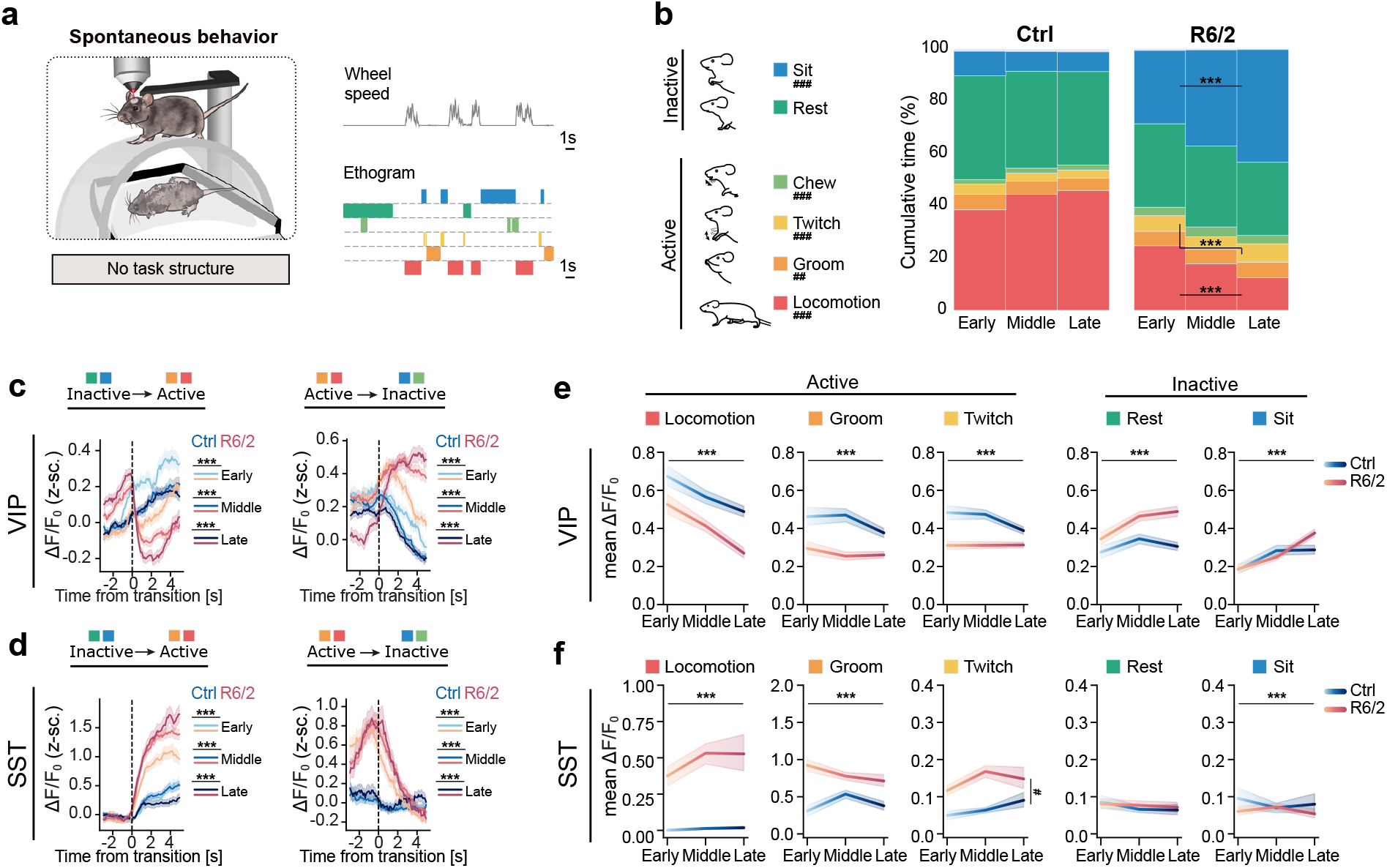
Abnormalities of spontaneous movements in HD mice are accompanied by subtype-specific dysfunctions of cortical INs. (**a**) Schematic of the experimental setup. Mice behaved on a passively rotating wheel with no task structure. *In vivo* calcium activity, wheel movement, and behavior ethogram were monitored simultaneously. (**b**) Cumulative duration of classified behavioral states. HD progression encompasses a progressive decrease of active periods in R6/2 mice (n = 26 ctrl and 24 R6/2 mice; Aligned rank transform; genotype main effect, locomotion *p* < 0.001, groom *p* = 0.008, twitch *p* < 0.001, chew *p* < 0.001, rest *p* = 0.463, sit *p* < 0.001; genotype:timepoint interactions: locomotion *p* < 0.001, twitch *p* < 0.001, sit *p* < 0.001, others n.s.). (**c**) Abnormal VIP-IN modulation in R6/2 mice at transitions from inactive to active behavioral states and vice versa. Average z-scored i1F/F0 aligned to state transitions (Aligned rank transform, fit separately for early, middle, and late timepoints; genotype:time bin interaction, *p* < 0.001 for all timepoints). (**d**) Same as (**c**), but for SST-INs (Aligned rank transform; genotype:time bin interaction, *p* < 0.001 for all timepoints). (**e**) Average VIP-IN activity is reduced during active behaviors in R6/2 mice (Aligned rank transform; genotype main factors: locomotion *p* = 0.021, groom *p* = 0.067, twitch *p* = 0.108, rest *p* = 0.048, sit *p* = 0.044; genotype:timepoint interactions: locomotion *p* < 0.001, groom *p* < 0.001, twitch *p* < 0.001, rest *p* < 0.001, sit *p* < 0.001). (**f**) Average SST-IN activity is increased during active behaviors in R6/2 mice (Aligned rank transform; genotype main factors: locomotion *p* = 0.080, groom *p* = 0.126, twitch *p* = 0.023, rest *p* = 0.782, sit *p* = 0.932; genotype:timepoint interactions: locomotion *p* < 0.001, groom *p* < 0.001, twitch *p* = 0.272, rest *p* = 0.412, sit *p* < 0.001). n = 8 VIP-Cre ctrl, n = 8 VIP-Cre::R6/2, n = 6 SST-Cre ctrl, n = 6 SST-Cre::R6/2 mice. Error bars represent mean ± c.i..

We proceeded to evaluate cortical IN activity during this behavioral paradigm in R6/2 and control mice. To achieve this, we expressed Cre-dependent GCaMP7f in M1 of VIP- and SST-Cre lines^17^, crossed to R6/2 mice and monitored VIP-IN and SST-IN activity longitudinally over 4-5 weeks, spanning early, middle, and late disease stages (**Extended Data Fig. 3a**). R6/2 mice displayed abnormal IN activity. In control mice, VIP-IN activity was higher during active behavioral states compared to inactive states. However, this modulation was reversed in R6/2 mice, in which VIP-INs became less active at the onset of active motor behaviors and more active at transitions to inactive behavioral states (**Fig. 2c, Extended Data Fig. 3b**). In contrast to VIP-INs, SST-INs in control mice showed only moderate activity during active behaviors, while SST-INs in R6/2 mice exhibited much more pronounced movement-related activity (**Fig. 2d, Extended Data Fig. 3c**). We evaluated IN alterations during episodes of all classified behaviors. In R6/2 mice, we observed a marked reduction in VIP-IN activity and increase in SST-IN activity during movements (locomotion, grooming and twitching), changes that were either absent or reversed during inactive states (rest and sit) (**Fig. 2e, f**).

**Fig. 3:**
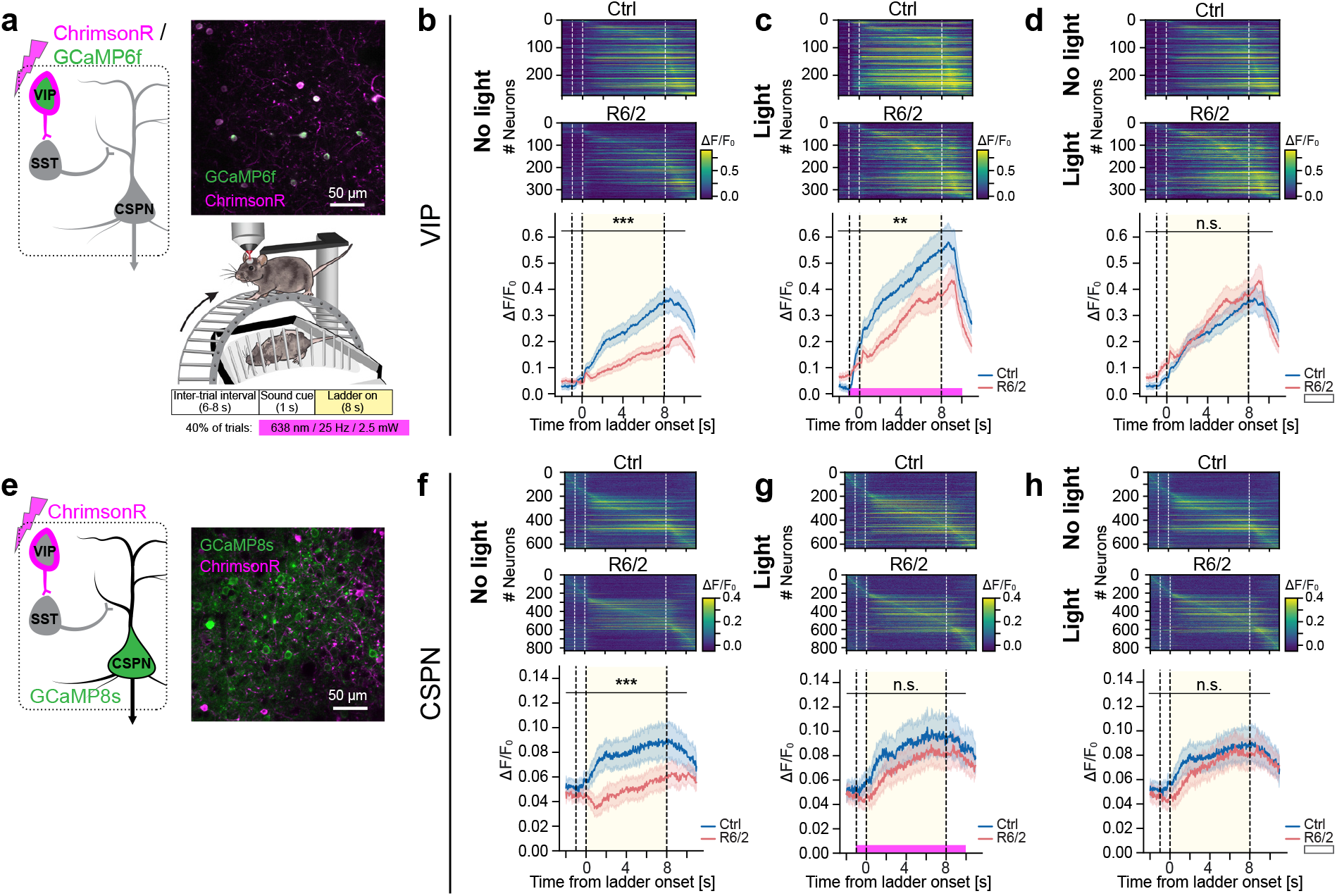
Optogenetic activation of VIP-INs normalizes VIP-IN and CSPN activity. (**a**) Left, scheme of VIP-IN stimulation and imaging. Upper right, example maximum intensity projections of *in vivo* two-photon fluorescence images in mice performing the ladder task, showing ChrimsonR and GCaMP6f co-expression in VIP-INs. Lower right, experimental setup and task structure. **b**) VIP-IN activity averaged over no-light trials (red masking light presented to the mouse). Dashed lines from left to right indicate: sound cue onset, ladder onset, and ladder offset. Data was binned into 2 s intervals for statistical comparison (n = 274 neurons / 9 sessions from 3 VIP-Cre ctrl mice and 343 neurons / 17 sessions from 5 VIP-Cre::R6/2 mice; Aligned rank transform; genotype main effect, *p* = 0.023; genotype:time bin interaction, *p* < 0.001). (**c**) Same as (**b**), but for light trials (638 nm, 25 Hz, 2.5 mW); (Aligned rank transform; genotype main effect, *p* = 0.167; genotype:time bin interaction, *p* = 0.002). (**d**) Comparison of no-light trials in control mice vs. light trials in R6/2 mice, replotted from (**b**) and (**c**); (Aligned rank transform; genotype main effect, *p* = 0.937; genotype:time bin interaction, *p* = 0.592). **(e)** Left, scheme of VIP-IN stimulation and CSPN imaging. Right, example maximum intensity projections of *in vivo* two-photon fluorescence images showing ChrimsonR expression in VIP-INs and GCaMP8s in CSPNs. (**f**) CSPN activity averaged over no-light trials (red masking light presented to the mouse). CSPNs showed reduced movement-related activation in R6/2 mice. (n = 636 neurons / 19 sessions from 5 VIP-Cre ctrl mice and 832 neurons / 33 sessions from 7 VIP-Cre::R6/2 mice, Aligned rank transform; genotype main effect, *p =* 0.182; genotype:time bin interaction, *p* < 0.001). (**g**) Same as (**f**), but for light trials (638 nm, 25 Hz, 2.5 mW). There is no difference in CSPN activity between ctrl and R6/2 mice (Aligned rank transform; genotype main effect, *p =* 0.810; genotype:time bin interaction, *p* = 0.867). (**h**) Comparison of no-light trials in control mice vs. light trials in R6/2 mice, replotted from (**f**) and (**g**); (Aligned rank transform; genotype main effect, *p* = 0.878; genotype:time bin interaction, *p* = 0.368). Error bars represent mean ± c.i.

Together, these results reveal that abnormal behavior during various types of spontaneous movements in R6/2 mice is accompanied by a consistent alteration in the activity of cortical neuron subtypes, with VIP-INs and SST-INs demonstrating hypoactivity and hyperactivity during movements, respectively.

### Optogenetic activation of VIP-INs restores CSPN activity in HD mice

The neuron subtype-specific alteration of activity in HD affords a potential opportunity for intervention to normalize cortical network function by a targeted modulation of genetically-defined neuron subtypes. VIP-INs are upstream of the other observed neuron types ^9–11,18^, and their hypoactivity in R6/2 mice could disinhibit SST-INs leading to their hyperactivity, which would in turn lead to increased inhibition of CSPNs explaining their hypoactivity. We hypothesized that artificial stimulation of VIP-INs could correct the hypoactivity of CSPNs in R6/2 mice. Therefore, we chose VIP-INs as the target for manipulation using optogenetics.

Neural activity modulation in diseases has an enormous therapeutic potential, but requires careful selection of the artificial modulation parameters to achieve physiological levels of neural activity^19^. To this end, we evaluated the effect of our VIP-IN stimulation protocol on VIP-IN activity associated with movement. We co-expressed the excitatory opsin ChrimsonR and GCaMP6f in VIP-INs in M1 of R6/2 mice to simultaneously activate them and image their activity. VIP-INs were activated with red light stimulation (638 nm, 25 Hz, 2.5 mW) in 40% of trials during the ladder task, starting at the auditory cue and terminating at 1 second after the ladder movement offset (**Fig. 3a, Extended Data Fig. 4a**). In the no-light trials, we observed a lower activity of VIP-INs in R6/2 mice during the movement period (**Fig. 3b**), consistent with the previous experiment (**Fig. 1i**). In the opto trials, VIP-INs were robustly activated, achieving overall activity levels in R6/2 mice similar to those seen in the no-light trials of control animals (**Fig. 3c, d**). Next, to test whether this normalization of VIP-IN activity can correct CSPN hypoactivity, we imaged CSPN activity while stimulating VIP-INs using the same protocol (**Fig. 3e, Extended Data Fig. 4b**). In the no-light trials, CSPNs showed hypoactivity during movements in R6/2 mice (**Fig. 3f**) as described above (**Fig. 1k**). In the opto trials, CSPN activity during movements of R6/2 mice was increased to the level statistically indistinguishable from that in no-light trials of control mice (**Fig. g, h**). Light stimulation alone did not have an effect on CSPN activity in control nor R6/2 mice in the absence of ChrimsonR (**Extended Data Fig. 4c-e**). These observations demonstrate that targeted VIP-IN activation can restore excitatory activity in downstream CSPNs and underscore the potential of circuit-level interventions to ameliorate network deficits in HD.

**Fig. 4:**
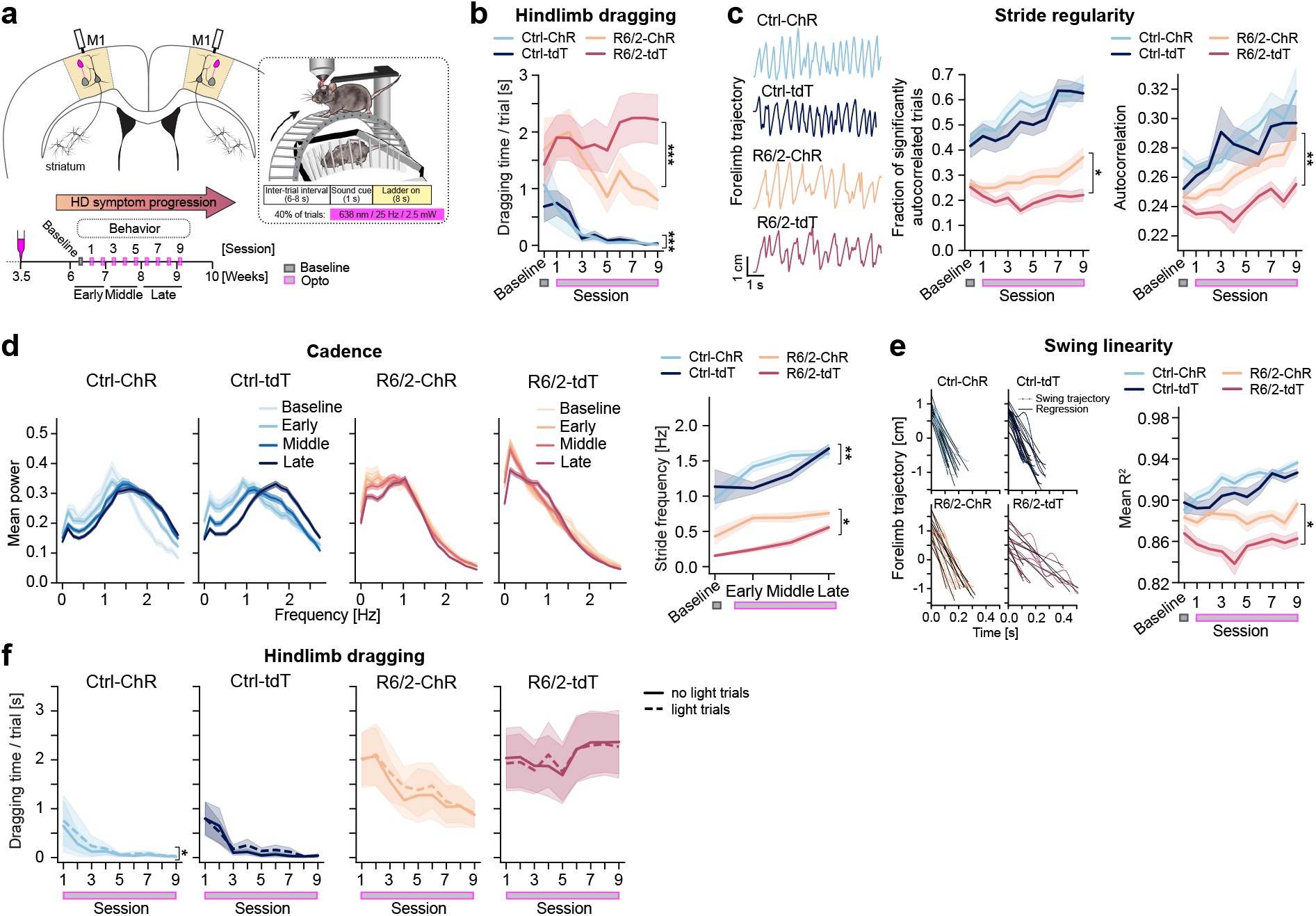
VIP-IN activation improves motor learning and ameliorates HD motor symptoms. (a) Schematic of the experimental paradigm and timeline. ChrimsonR was expressed bilaterally in M1 VIP-INs, and light dispersing cannulas were implanted over the cortical surface. Following two training sessions and one baseline recording, mice performed in the ladder task paired with optogenetic stimulation in 40% of trials every other day for 9 sessions. (b) Duration of hindlimb dragging per trial decreased in ctrl-ChR, ctrl-tdTomato, and R6/2-ChR mice (Aligned rank transform; actuator:session number interaction, ctrl: *p* < 0.001, R6/2: *p* < 0.001). (**c**) Left, example movement trajectories of the left forelimb in single trials at a late timepoint. Right, autocorrelation performed on forelimb trajectories in individual trials reveals that the regularity of oscillatory gait pattern increased with learning in ctrl-ChR, ctrl-tdTomato, and R6/2-ChR mice: fraction of significantly autocorrelated trials and max. autocorrelation value per trial (Aligned rank transform; actuator:session number interaction; fraction of significantly autocorrelated trials; ctrl: *p* = 0.492, R6/2: *p* = 0.044; autocorrelation; ctrl: *p* = 0.727, R6/2 *p* = 0.018). (**d**) Left, trial-averaged power spectra of forelimb trajectories. With motor learning, cadence increased and low frequency components of gait decreased in ctrl-ChR and ctrl-tdTomato mice. R6/2-ChR mice showed a shift in the average spectrum resembling that of ctrl mice. Right, mean peak frequencies of the plots on the left. Cadence increased in R6/2-ChR mice over time (Aligned rank transform; actuator:session number interaction, ctrl: *p* = 0.002, R6/2: *p* = 0.032). (**e**) Regression analysis of individual forelimb strides. Left: strides of single example trials, aligned to the swing onset of the stride phase, overlaid with regression fits. Right: average R^2^ values – representing a measure for the directness of swings – increased with motor learning and VIP-IN stimulation in ctrl and R6/2 mice (Aligned rank transform; actuator:session number interaction, ctrl: *p* = 0.146, R6/2: *p* = 0.020). (**f**) The effects of VIP-IN stimulation in R6/2 mice last beyond the light trials. Comparison of no-light and light trials for hindlimb dragging durations (Aligned-rank transform; trial type:session number interaction, *p* = 0.016 for Ctrl-ChR, *p* = 0.809 for Ctrl-tdTomato, *p* = 0.999 for R6/2-ChR, *p* = 0.999 for R6/2-tdTomato). n = 9 VIP-Cre-ChR, n = 10 VIP-Cre-tdT, 11 VIP-Cre::R6/2-ChR, and 11 VIP-Cre::R6/2-tdT mice, 779 sessions across all mice and timepoints. Error bars represent mean ± s.e.m.

### VIP-IN activation improves motor learning and ameliorates motor symptoms in HD mice

Does the restoration of cortical network activity improve motor behavior in R6/2 mice? To address this question, we used the stimulation protocol to activate VIP-INs and examined its effects on motor performance in R6/2 mice. We expressed ChrimsonR bilaterally in the M1 cortex (**Extended Data Fig. 5a, Methods**). At six weeks of age, corresponding to an early disease stage, mice completed a baseline session on the ladder task without optogenetic stimulation. Following this, mice were subjected to the ladder task every other day across nine sessions, with 40% of randomly interspersed trials with red light stimulation bilaterally (638 nm, 25 Hz, 2.5 mW) (**Fig. 4a**). Across sessions, control mice improved their task performance, demonstrated by reduced hindlimb dragging, increased autocorrelation of gait trajectories, a clear peak in cadence rhythm, and increased regularity of strides (**Fig. 4b-e, Extended Data Fig. 5b**). Consistent with the previous results (**Fig. 1d-g**), R6/2 mice expressing tdTomato as a control showed persistent hindlimb dragging without improvement, irregular gait patterns without a coherent cadence rhythm, and poor coordination of strides (**Fig. 4b-e, Extended Data Fig. 5b**). In contrast, R6/2 mice that received VIP-IN stimulation displayed a progressive decrease in hindlimb dragging over time (**Fig. 4b**), and more regular stride patterns compared to R6/2 tdTomato mice (**Fig. 4c**). Furthermore, R6/2 mice receiving VIP-IN stimulation demonstrated cadence changes similar to control mice, adopting faster and more consistent strides over time (**Fig. 4d**), as well as substantial improvements in the directness of stride swings (**Fig. 4e**).

In the analysis above, the light trials (40%) and no-light trials (60%) were pooled together. We next sought to examine whether the beneficial effects of VIP-IN stimulation were only during the stimulation, or whether they persisted beyond stimulation. To this goal, we compared the behavioral performance in light and no-light trials. Strikingly, we did not find any difference in behavioral performance between these trial types in R6/2 mice with VIP-IN stimulation (**Fig. 4f, Extended Data Fig. 5b**). These results suggest that VIP-IN stimulation during movements have lasting benefits, likely by promoting motor learning.

These experiments demonstrate that targeted VIP-IN activation can mitigate motor deficits in HD.

## Discussion

The symptoms of neurological disorders arise from a complex interplay of genetic, molecular, and circuit-level dysfunctions, making it difficult to design therapeutics to ameliorate the symptoms. This is true even in HD, which is a monogenic disorder with a clearly defined genetic origin. Identifying the critical nodes of the complex dysfunctions would illuminate intervention strategies, as modulation of these nodes could correct multiple downstream processes.

Cortical dysfunction in HD is among the earliest pathological signatures, which includes an imbalance between excitation and inhibition^20–22^. Studies in conditional HD mouse models have found that mHTT expression in cortical INs and principal neurons both contribute to the development of full-fledged cortical HD pathology and behavioral defects ^8,23^. Furthermore, CSPNs in the motor cortex appear selectively vulnerable in HD^24^. While previous work suggests differential contributions of cortical neuron subtypes to driving behavioral symptoms^21,25,26^, the roles of specific inhibitory and excitatory neuron populations remain poorly understood.

Here, we examined the activity patterns of two cortical IN subtypes and CSPNs in the well-established R6/2 HD mouse model^13^ *in vivo* and during identified behaviors. We observed neuron subtype-specific abnormalities in their activity which were consistent across learned and spontaneous movements. Of these abnormalities, the hypoactivity of VIP-INs appears to be an upstream node, as selective restoration of VIP-IN activity with optogenetics in R6/2 mice corrected the hypoactivity of CSPNs during movement. Remarkably, this intervention within the motor cortex improved motor deficits over time. VIP-INs are a unique group of cortical INs with preferential connections onto other INs and thus their activity results in disinhibition of the excitatory population. Even though our stimulation parameters were carefully titrated, it is still striking that the artificial pattern of stimulation benefited behavior, considering that behavior is controlled by precise and complex spatiotemporal patterns of neural activity. We speculate that our manipulation on the inhibitory circuits effectively reduced the pathologically strong levels of inhibition on excitatory neurons, such that excitatory neurons could now function more normally to express their healthy activity patterns.

VIP-IN activity is sensitive to behavioral states and, through their disinhibitory function, these cells contribute to the modulation of cortical gain and plasticity ^10,27–29^. Consistently with their role in regulating plasticity, we found that the beneficial effects of VIP-IN stimulation carried over to the subsequent trials without VIP-IN stimulation. This observation suggests that the activation of VIP-INs does not only facilitate activity patterns related to movement execution, but also ‘ungates’ synaptic plasticity mechanisms required for motor learning. The lasting benefits of VIP-IN stimulation suggest that the potential therapeutic applications of this approach may not require a constant stimulation over long periods of time.

An endogenous mechanism mediating VIP-IN activity regulation involves cholinergic input from basal forebrain projections, which are known to drive VIP-IN-mediated disinhibition during motor learning^10^. Cholinergic signaling is essential to facilitate synaptic plasticity and motor adaptation, processes that rely on VIP-IN-mediated modulation of excitatory output. In HD, early cholinergic deficits^30,31^ may disrupt this modulatory pathway, potentially contributing to VIP-IN dysfunction.

Our results suggest that VIP-INs may be a critical target for intervention in HD. Their strikingly low-dimensional activity changes seen in our experiments make them feasible targets for directional activity manipulation. Additionally, the fact that they show relatively slow accumulation of intranuclear mHTT inclusion bodies and possibly are less vulnerable to cell-autonomous toxicity mechanisms than principal neurons^32^ may further argue for their suitability as a therapeutic target. More broadly, our results underscore the therapeutic potential of targeting specific neuron subtypes to rebalance neural circuit dynamics. This approach could inform future therapies aimed at modulating the activity of distinct neuronal populations to alleviate symptoms and slow disease progression in HD and potentially other neurological disorders with distinct circuit-level disruptions. It is important to note, however, that this type of circuit-based intervention likely would not stop neurodegeneration. Thus, the circuit rebalancing approach for behavioral benefits should be pursued in parallel with approaches to target the root causes of neurodegeneration.

## Acknowledgments

We thank B. Morales, A. Medina, M. Boehm and E. Hall for technical assistance; S. de Vries for insightful discussion on cortical neuron types; G. Mishne, M. Aoi, P. Goltstein and S. Reinert for helpful advice on analyses; R.D. Sristi for code used for preprocessing; all members of the Komiyama and Dudanova labs, especially N. G. Hedrick and K.Völkl for helpful discussions on the project and manuscript.

## Funding

The work was supported by the following funding sources:

National Institutes of Health grant R01 NS125298 (T.K.)

National Institutes of Health grant R01 NS091010 (T.K.)

National Institutes of Health grant R01 DC018545 (T.K.)

National Institutes of Health grant R01 MH128746 (T.K.)

National Science Foundation grant 2024776 (T.K.)

Simons Collaboration on the Global Brain Pilot Award (T.K.)

Max Planck Society for the Advancement of Science (R.K. and I.D.)

Deutsche Forschungsgemeinschaft (DFG, German Research Foundation), SFB1451 project number 431549029-A09 (I.D.)

Deutsche Forschungsgemeinschaft, Germany’s Excellence Strategy – CECAD, EXC 2030-390661388 (I.D.)

Chan-Zuckerberg Initiative (T.K. and I.D.)

European Research Council (ERC) Synergy Grant under FP7 GA number ERC-2012-*SyG_318987*-Toxic Protein Aggregation in Neurodegeneration (ToPAG) (R.K.)

DFG Walter Benjamin Postdoctoral Fellowship and DFG Return Fellowship BL1812 (S.B.) Joachim Herz Foundation Add-On Fellowship for Interdisciplinary Life Science (S.B.)

## Author contributions

Conceptualization: SB, ID, TK

Methodology: SB, EG, TK

Behavioral training and histology: SB, YS, SS

Investigation: SB, DA

Visualization: SB

Formal analysis: SB

Software: SB, NDG

Data curation: SB, NDG

Funding acquisition: SB, ID, RK, TK

Project administration: ID, TK

Supervision: ID, RK, TK

Writing – original draft: SB, TK

Writing – review & editing: SB, ID, TK with input from all authors

## Data and materials availability

All data needed to evaluate the conclusions in the paper are present in the paper and/or Supplemental Materials. Data and code used to analyze data and generate figures for this paper are available upon request from the corresponding authors.

## Supplementary Materials

Materials and Methods

Extended Data Figs. 1-5

Extended Data Table 1

References (*33-44*)

## References

1. Arber, S. & Costa, R. M. Connecting neuronal circuits for movement. Science 360, 1403–1404 (2018).

2. The Huntington’s Disease Collaborative Research Group. A novel gene containing a trinucleotide repeat that is expanded and unstable on Huntington’s disease chromosomes. Cell 72, 971–983 (1993).

3. Shepherd, G. M. G. Corticostriatal connectivity and its role in disease. Nat Rev Neurosci 14, 278–291 (2013).

4. Bunner, K. D. & Rebec, G. V. Corticostriatal Dysfunction in Huntington’s Disease: The Basics. Front Hum Neurosci 10, 317 (2016).

5. Fernández-García, S. et al. M2 cortex-dorsolateral striatum stimulation reverses motor symptoms and synaptic deficits in Huntington’s disease. Elife 9, e57017 (2020).

6. Kim, E. H. et al. Cortical interneuron loss and symptom heterogeneity in Huntington disease: Interneuron Loss in HD Cortex. Ann Neurol 75, 717–727 (2014).

7. Philpott, A. L. et al. Cortical inhibitory deficits in premanifest and early Huntington’s disease. Behav Brain Res 296, 311–317 (2016).

8. Gu, X. et al. Pathological Cell-Cell Interactions Elicited by a Neuropathogenic Form of Mutant Huntingtin Contribute to Cortical Pathogenesis in HD Mice. Neuron 46, 433–444 (2005).

9. Pfeffer, C. K., Xue, M., He, M., Huang, Z. J. & Scanziani, M. Inhibition of inhibition in visual cortex: the logic of connections between molecularly distinct interneurons. Nat Neurosci 16, 1068–1076 (2013).

10. Ren, C. et al. Global and subtype-specific modulation of cortical inhibitory neurons regulated by acetylcholine during motor learning. Neuron (2022) doi:10.1016/j.neuron.2022.04.031.

11. Kepecs, A. & Fishell, G. Interneuron cell types are fit to function. Nature 505, 318–326 (2014).

12. Jackson, J., Ayzenshtat, I., Karnani, M. M. & Yuste, R. VIP+ interneurons control neocortical activity across brain states. J Neurophysiol 115, 3008–3017 (2016).

13. Mangiarini, L. et al. Exon 1 of the HD Gene with an Expanded CAG Repeat Is Sufficient to Cause a Progressive Neurological Phenotype in Transgenic Mice. Cell 87, 493–506 (1996).

14. Mathis, A. et al. DeepLabCut: markerless pose estimation of user-defined body parts with deep learning. Nat Neurosci 21, 1281–1289 (2018).

15. Taniguchi, H. et al. A Resource of Cre Driver Lines for Genetic Targeting of GABAergic Neurons in Cerebral Cortex. Neuron 71, 995–1013 (2011).

16. Bohnslav, J. P. et al. DeepEthogram, a machine learning pipeline for supervised behavior classification from raw pixels. Elife 10, e63377 (2021).

17. Hippenmeyer, S. et al. A Developmental Switch in the Response of DRG Neurons to ETS Transcription Factor Signaling. PLoS Biol. 3, e159 (2005).

18. Millman, D. J. et al. VIP interneurons in mouse primary visual cortex selectively enhance responses to weak but specific stimuli. Elife 9, e55130 (2020).

19. Hattori, R., Kuchibhotla, K. V., Froemke, R. C. & Komiyama, T. Functions and dysfunctions of neocortical inhibitory neuron subtypes. Nat Neurosci 20, 1199–1208 (2017).

20. Cummings, D. M. et al. Alterations in Cortical Excitation and Inhibition in Genetic Mouse Models of Huntington’s Disease. J Neurosci 29, 10371–10386 (2009).

21. Burgold, J. et al. Cortical circuit alterations precede motor impairments in Huntington’s disease mice. Sci Rep-uk 9, 6634 (2019).

22. Spampanato, J., Gu, X., Yang, X. W. & Mody, I. Progressive synaptic pathology of motor cortical neurons in a BAC transgenic mouse model of Huntington’s disease. Neuroscience 157, 606–620 (2008).

23. Dougherty, S. E. et al. Hyperactivity and cortical disinhibition in mice with restricted expression of mutant huntingtin to parvalbumin-positive cells. Neurobiol Dis 62, 160–171 (2014).

24. Pressl, C. et al. Selective vulnerability of layer 5a corticostriatal neurons in Huntington’s disease. Neuron 112, 924-941.e10 (2024).

25. Arnoux, I. et al. Metformin reverses early cortical network dysfunction and behavior changes in Huntington’s disease. Elife 7, e38744 (2018).

26. Donzis, E. J. et al. Cortical Network Dynamics Is Altered in Mouse Models of Huntington’s Disease. Cereb Cortex 30, 2372–2388 (2019).

27. Fu, Y. et al. A Cortical Circuit for Gain Control by Behavioral State. Cell 156, 1139–1152 (2014).

28. Fu, Y., Kaneko, M., Tang, Y., Alvarez-Buylla, A. & Stryker, M. P. A cortical disinhibitory circuit for enhancing adult plasticity. Elife 4, e05558 (2015).

29. Pi, H.-J. et al. Cortical interneurons that specialize in disinhibitory control. Nature 503, 521–524 (2013).

30. Smith, R. et al. Cholinergic neuronal defect without cell loss in Huntington’s disease. Hum. Mol. Genet. 15, 3119–3131 (2006).

31. Pancani, T. et al. Cholinergic deficits selectively boost cortical intratelencephalic control of striatum in male Huntington’s disease model mice. Nat Commun 14, 1398 (2023).

32. Voelkl, K., Schulz-Trieglaff, E. K., Klein, R. & Dudanova, I. Distinct histological alterations of cortical interneuron types in mouse models of Huntington’s disease. Front Neurosci-switz 16, 1022251 (2022).

