## Supplemental Information for "Optogenetic restoration of neuron subtype-specific cortical activity ameliorates motor deficits in Huntington’s Disease mice"

### Materials and Methods

#### *Animals*

All animal procedures were performed in accordance with guidelines set forth and protocols approved by the UCSD Institutional Animal Care and Use Committee and the National Institutes of Health, as well as by the Government of Upper Bavaria, Germany (animal protocols 55.2-1-54-2532-168-2014, 55.2-1-54-2532-19-2015, 55.2-2532.Vet_02-20-05, and 55.2-2532.Vet_02-19-83). R6/2 mice[1] transgenic for the 5′ end of the human huntingtin gene were obtained from Jackson Laboratories and maintained by crossing R6/2 males to F1 C57Bl6/CBA females. The presence of the transgene was verified by PCR with the following primers: forward, 5′CCGCTCAGGTTCTGCTTTTA-3′, reverse, 5′-TGGAAGGACTTGAGGGACTC-3′. CAG repeat length was determined by Laragen for all experimental groups. Spontaneous behavior experiments on the wheel were performed at the Max Planck Institute for Biological Intelligence, and motorized ladder experiments were performed at UCSD. Separate batches of R6/2 mice were used for these two sets of experiments, and the CAG repeat lengths were different between these two groups (202 ± 13 and 157 ± 6 for Max Planck Institute and UCSD respectively, mean ± SD) which led to different speeds of disease progression. Therefore, the stages of disease progression were matched across these batches of mice by monitoring their body weights (**Extended Data Fig. 1a and 2a**). Mice were group housed in cages with standard bedding in a temperature-controlled room (~21ºC) with a reversed 12hr light/12hr dark cycle. Mice were allowed *ad libitum* access to food and water. Both male and female mice were used for all experiments.

#### *Surgery*

Surgical procedures were performed as previously described [2–4]. Briefly, 3.5-week-old mice were anaesthetized with an intraperitoneal (i.p.) injection of ketamine/xylazine (130 and 8 mg $kg^{-1}$ body weight, respectively) and a low dose of isoflurane (0.5 % with constant flow rate of 1L $min^{-1}$ at 0.1 Bar). After reaching a deep plane of anesthesia, Baytril (10 mg/kg) and dexamethasone (5 mg/kg) were injected subcutaneously to prevent infection and brain swelling, respectively.

For imaging experiments, a craniotomy (4 mm diameter) was performed, as previously described[2–4], over the right caudal forelimb area of M1 centered at 0.5 mm anterior and 1.5 mm lateral from bregma. For imaging and manipulating cortical inhibitory neurons, viruses (calcium sensors: AAV9-hSyn-FLEX-jGCaMP7f or AAV2/1-hSyn-FLEX-GCaMP6f, titer: ~$10^{12}$ vg/ml; opsin / control: AAV5-Syn-FLEX-rc[ChrimsonR-tdTomato] or AAV2/1-CAG-FLEX-tdTomato, titer: ~$10^{13}$ vg/ml; Addgene) were injected into the caudal forelimb area of M1 using a beveled glass pipette (~12-25 µm inner diameter). Each injection consisted of a ~200 nl volume centered at a depth of ~400 µm below the pial surface. 3 injections, separated by at least 500 µm horizontally, were performed in each craniotomy.

For imaging corticostriatal projection neurons, retrograde virus (rgAAV-hSyn-jGCaMP8s, titer ~$10^{13}$ infecting units per ml, Addgene) was injected into the dorsolateral striatum at 0.5 mm anterior, 3.17 mm lateral from bregma. Two injections, each consisting of a ~300 nl volume were performed at a 22° angle and at 2.2 mm and 2.0 mm below the pial surface, respectively. After virus injections, the glass pipette was left in place for 5 minutes to avoid backflow. Following injections, a round coverslip (VWR; d = 4 mm) was implanted into the craniotomy and affixed to the skull using histoacryl glue (B.Braun) and dental acrylic cement.

For longitudinal optogenetic stimulation experiments, viruses (AAV5-Syn-FLEX-rc[ChrimsonR-tdTomato] or AAV2/1-CAG-FLEX-tdTomato, titer: ~$10^{13}$ vg/ml; Addgene) were injected at two sites per hemisphere through small burr holes. Each injection consisted of a ~200 nl volume, injection sites were at +0.5 AP/± 1.5 ML and +2.0 AP/±1.3 ML. Fiber optic cannulas (Doric, 600 µm core diameter / 0.22 NA, diffuser tip) were implanted at a 15° angle onto the cortical surface of both hemispheres at +1.0 AP/± 1.0 ML. A custom-built head-bar was glued and cemented to the skull to allow for stable head fixation. An analgesic (Buprenorphine (0.1 mg/kg) or Carprofen (5 mg/kg)) was injected ~1 hr prior to the end of the surgery to manage postoperative pain. Following surgery, mice were administered daily Baytril, dexamethasone, and analgesic for up to 3 days to manage postoperative infection, swelling, and pain, respectively.

#### *Behavior*

*Handling and training*

At the age of 5 weeks, mice were handled on 4 consecutive days for 10 min until they were familiarized with the trainer and routinely ran from hand to hand. In the subsequent behavior task training, mice got adjusted to the experimental setup and head fixation. Training sessions (30 min each, 4 consecutive days) were performed in the dark with an IR light source for the camera.

*Motorized ladder task*

Head-fixed mice were positioned onto a custom-built, circular ladder (diameter = 19 cm, rung spacing = 1 cm), adapted from the KineMouse Wheel (https://hackaday.io/project/160744-kinemouse-wheel) and motorized by an electric DC motor (12V, 60 RPM). Trial structure was controlled using BPod (v0.5). In each trial, 8 seconds of ladder rotation (speed ~ 10 mm/s) were preceded by an auditory cue (1 second at 3 kHz) for 1 second. Trials were separated by a variable 6-8 second inter-trial interval. In each trial, the auditory cue (= trial start) was indicated by a ~30 ms IR LED flash, facilitating the alignment of trials during video analysis.

*Wheel task*

Head-fixed mice were placed on a passively rotating wheel (KineMouse Wheel, d = 19 cm, continuous surface)[5]. No cues or trial structure were provided and mice typically showed alternating active and inactive behavioral states. Running wheel motion was captured by a rotary encoder (1000 CPR, 60 Hz sampling rate). Speed and directionality of the wheel were decoded online using a Tensy 3.2 processor board custom written Arduino code, adapted from Janelia Open Science Laboratory Tools (https://www.janelia.org/open-science/encoder-interface-for-mouse-treadmill).

*Behavioral videography*

For both behavioral tasks, two orthogonal views of the mouse were captured simultaneously through a mirror mounted at 45˚ inside the ladder and wheel. Mouse behavior was tracked at 60 - 100 Hz with an IR-sensitive video camera (USB 2.0, 1/3″CMOS, 744 × 480 pixel; 8 mm M0814MP2 1.4–16 C, 2/3″, megapixel c-mount objective; The Imaging Source) and custom software (Input Controller, The Imaging Source).

#### *In vivo two-photon imaging*

Imaging during the motorized ladder task was performed using a commercial two-photon microscope (MOM, Sutter Instrument) equipped with a ×16/0.8-NA objective (Nikon) and a Ti:Sapphire excitation laser (Mai Tai, Spectra-Physics) tuned to 925 nm. Images (512 × 512 pixels, ~1 µm/px) were recorded at ~29 Hz for the duration of the behavioral session (~12 min,

40-50 trials). Frame times were recorded and synchronized with behavioral recordings (videography).

Imaging during the wheel task was performed using a commercial two-photon microscope (B-Scope, ThorLabs) equipped with a ×16/0.8-NA objective (Nikon) and a InsightDS+ laser (Spectra-Physics) tuned to 925 nm. Image acquisition was controlled through ThorImage 4.0 software. Images (768 × 768 pixels, 0.702 µm/px) were recorded at ~10 Hz for the duration of the behavioral session (10 min per field of view (FOV), 4-5 FOVs per session). Frame times were recorded and synchronized with behavioral recordings (videography, rotary encoder) using the ThorSync software.

Laser power at the objective was controlled with a Pockel's cell (Conoptics) and ranged between 10-50 mW for all experiments. Depth of FOVs below the cortical surface was between 120 – 330 µm for VIP-INs, 120 – 550 µm for SST-INs and 200 – 540 µm for CSPNs. For longitudinal repositioning, an epifluorescent image was acquired in the first imaging session to capture vasculature, and xyz-coordinates provided by the microscope stage were documented for each FOV. The average projection two-photon image of 100 frames was used as a reference in later sessions and the z-plane was carefully adjusted to maximally match the imaging plane to the reference image.

#### *Optogenetics*

For simultaneous *in vivo* two-photon imaging and photoactivation of VIP-INs, light from a red diode laser (Oxxius LBX-638-HPE, 638 nm) was delivered through the objective of the microscope. To avoid interference between stimulation and imaging, the optogenetic light was synchronized with the resonant scanner, delivering a pulse of light at the turnaround of the scanner (20 µs for each period) at 2.5 mW power and an effective frequency of 25 Hz. For optogenetic behavior experiments (50 trials / session), red laser light (Oxxius LBX-638-HPE, 638 nm) was delivered through a bifurcated silica fiber optic patch cord (Doric, 400 µm core diameter / 0.22 NA) at 25 Hz / 2.5 mW and through implanted light diffusing fiber optic cannulas (Doric, 600 µm core diameter / 0.22 NA, diffuser tip) implanted over M1 of both hemispheres. In optogenetic experiments, to avoid visual effects of the stimulation light, mice were presented with a red masking LED light in all trials.

#### *Image analysis*

A combination of Suite2P and Cellpose[6,7] software was used to generate regions of interest (ROIs) corresponding to individual neurons and to extract their fluorescence. ROI classifications by the automatic classifier were further refined by manual inspection. A fluorescence trace's time-varying baseline ($F_0$) was estimated using custom-written MATLAB code by smoothing inactive portions of the trace using a previously described iterative procedure[5]. Briefly, this process identified the trace's active and inactive portions, removing active portions and using the LOESS-smoothed inactive portions (interpolated across active periods) to estimate the time-varying baseline. The normalized $\Delta F/F_0$ trace was then calculated, where $\Delta F$ was found by subtracting the baseline trace from the raw trace, and $F_0$ was the calculated time-varying baseline. A calcium activity event trace was constructed which was zero except for frames with detected events, as previously described [2].

#### *Behavior classification*

For deep-learning assisted classification of innate behaviors from raw videos, we used DeepEthogram (DEG)[8]. A single DEG model was trained in an iterative fashion. Based on our visual observations of mouse behavior, we defined a set of behavior classes of interest: 'locomotion', 'grooming', 'rest' (an inactive state with both forelimbs resting on the wheel surface), 'sit' (an inactive state with both forelimbs resting above the wheel surface), 'twitch' (short, fast forelimb twitches) and 'chew' (jaw movements, including licking). An initial model was trained on ~10,000 manually labeled frames from a total of six exemplary videos of control and R6/2 mice of different ages. Model predictions of the initial and a few additional videos were corrected manually and used for retraining the model for the next iteration. A total of 19 training iterations was performed on a final number of 66 videos, until all behavior classes of interest were detected with high F1 scores > 0.7. The model also identified 'shake' (a full-body twitching behavior), which occurred in 1.1% of frames. However, 'shake' almost always occurred concurrently with other behavior classes and was excluded from further analysis.

*Voluntary behavior-related activity*

To estimate the activity of individual neurons at the transition between active and inactive behaviors, we only considered neurons with a non-zero number of calcium events detected in a given session. Each neuron's activity (z-scored $\Delta F/F_0$) was aligned to the transition between behavior classes. We considered 'locomotion' and 'grooming', classified using DEG as active motor behaviors, while 'rest' and 'sit' classes jointly represented the inactive behavioral periods. We only considered transitions between classes that were at least 3 seconds long to avoid contamination of activity related to other behaviors. 'Chew' and 'twitch' generally did not meet this criterium and their transitions were not considered for this analysis. Each neuron's activity was averaged across all individual transitions. For analysis of population activity during identified behavior classes, we averaged each neurons activity ($\Delta F/F_0$) across all episodes of a classified behavior. We did not consider 'Chew' for this analysis, as it was rare (2.6% of frames) and almost never occurred in isolation without another concurrent behavior (0.005% of frames).

### ***Gait analysis***

*Data preprocessing*

We used Python 3.8 and relevant libraries to process and analyze behavior data. To extract limb movement trajectories from behavior videos, we used DeepLabCut (DLC) [9]. A single DLC model was trained using video frames from across a sample of experiments. Limb movements were captured from two perspectives (side and bottom). Optical distortion through the mirror was corrected by aligning x-coordinates of side and bottom perspectives. Corrected x-coordinates from both perspectives, detected at >95 % DLC likelihood, were averaged. Trial information captured with BPod (v0.5) was aligned and experimental metadata was added to each session. To preserve the essential motion characteristics while filtering out high-frequency noise, we used a Butterworth filter with a normalized cutoff removing frequencies from paw trajectories that were less than one-tenth of the Nyquist frequency (i.e., half the video frame rate).

*Hindlimb dragging analysis*

We calculated the average Euclidian distance from left forelimb to right hindlimb and from right forelimb to left hindlimb and normalized the distance by the mouse body weight[10]. To calculate the time spent dragging, the cumulative time for which the normalized forelimb - hindlimb distance exceeded 2 standard deviations above the control population mean was calculated for each trial (Extended Data Fig. 1c).

*Autocorrelation analysis*

To analyze regularity in gait patterns, we autocorrelated limb trajectories in each individual ladder trial. To this end, we centered the trajectories around zero for each trial, removing offsets. We used the statsmodels ACF package to compute the autocorrelation function for the x-position of each limb, measuring self-similarity over lags up to 5 seconds in each 8 second trial. The maximum peak of the autocorrelation function was determined. A peak was considered significant if it exceeded the upper bound of the 95% confidence interval, indicating significantly rhythmic behavior in the respective trial. The fraction of significantly autocorrelated trials was determined by dividing the number of significant trials by the total number of trials.

*Fourier analysis*

To capture movement frequencies during the gait cycle, we processed the x-coordinate trajectories of forelimbs through Fourier transformation. In short, we converted the z-scored trajectory of each forelimb in each trial from the time domain to the frequency domain using the scipy fft function. The power spectrum was then computed as the squared magnitude of the Fourier coefficients, representing the energy associated with each frequency component. Average power spectra were obtained by calculating the mean power within 0.13 Hz bins. The power spectrum was normalized by dividing each binned power by the maximum power, scaling the values to a range between 0 and 1, allowing for comparison across sessions and animals. The maximum of the normalized power spectrum was used to extract the dominant stride frequency per session and animal.

*Linear regression analysis*

To analyze the directness of strides, we used linear regression on isolated stride phases. The x-coordinates of zero-centered forelimb trajectories of individual trials were segmented into swing and stride phases using the peakdetect library. We defined a minimum difference in the signal required to identify extrema in the trace (5 mm). Based on these extrema points, the trace was segmented and labeled ‘swing’ and ‘stance’ depending on whether the trace was rising or falling between extrema, respectively. We used linear regression on each swing trajectory to determine the $R^2$ score, measuring the ‘directness’ of the trajectory.

***Histology***

Mice were transcardially perfused with 0.1M sodium phosphate buffered saline (PBS) followed by 4% paraformaldehyde (PFA, wt/wt) in 0.1M PBS. Brains were removed and postfixed in 4% PFA overnight at 4ºC. Following postfix, whole brains were stored in 30% sucrose in 0.1M PBS at 4ºC for 2-3 days. Coronal sections (50 µm thick) containing the motor cortex were then prepared using a Leica SM 2000R sliding microtome. Floating sections were washed in 0.1M PBS 3×5 min. mounted on slides using CC Mount (Sigma-Aldrich) and allowed to cure overnight before imaging.

***Statistics***

Statistical tests were selected based on data distributions and significance was set at $p < 0.05$. For neuronal activity analysis, we averaged a neuron’s activity over trials or behavioral epochs in a given session. Longitudinal behavioral measures were averaged on the session level. All evaluated datasets were tested for normality and non-parametric tests were used throughout the manuscript where appropriate. No statistical methods were used to pre-determine sample sizes, but our sample sizes are similar to those reported in previous publications[2,4].

To compare two or more groups (e.g. genotypes, disease stage, time before and after behavior onset), mixed-effects models were used to account for the nested structure of the data and minimize effects introduced by inter-animal variability[11,12]. For statistical estimation of non-normally distributed datasets, we used an Aligned Rank Transform (ART) variant of mixed-effects models using the ARTools packaged from R, which allows for nonparametric mixed-effects model testing. Multiple comparisons were corrected using the two-stage False Discovery Rate (FDR) method.

Two-factor mixed-effects models for estimating mouse body weight across disease progression (Extended Data Fig. 1a, 2a) were constructed as follows:

$$y \sim genotype + week + genotype{:}week + (1\ |animal)$$

with fixed main effect terms for group identity (ctrl, R6/2) and mouse age in weeks, a fixed interaction term for the interaction between group identity and week, and a random effect term grouped by animal. If a significant interaction was detected, posthoc pairwise Mann-Whitney U tests were performed between groups for each week.

Two-factor mixed-effects models for estimating behavior performance and activity levels across disease progression (Figures 1d, 1e, 1g, 2e, 2f, Extended Data Fig. 1d, 1e) were constructed as follows:

$$y \sim genotype + timepoint + genotype{:}timepoint + (1\ |animal)$$

with fixed main effect terms for group identity (ctrl, R6/2) and disease stage (early, middle, late), a fixed interaction term for the interaction between group identity and disease stage, and a random effect term grouped by animal. If a significant interaction was detected, posthoc pairwise Mann-Whitney U tests were performed between groups for each disease stage.

Similarly, to compare population average activity aligned to ladder task trials or behavior transitions (Figures 1i, 1j, 1k, 2c, 2d, 3b, 3c, 3d, 3f, 3g, 3h, Extended Data Fig. 4d, 4e), the models were constructed as follows:

$$y \sim genotype + time\ bin + genotype{:}time\ bin + (1\ |animal)$$

with fixed main effect terms for genotype (ctrl, R6/2) and 2-second time bins (-2-0, 0-2, 2-4, 4-6, 6-8, 8-10, 10-12 seconds relative to ladder onset or -2-0, 0-2, 2-4 seconds relative to behavior transition), the interaction term between genotype and time bin, and a random effect term grouped by animal. If a significant interaction was detected, posthoc pairwise Mann-Whitney U tests were performed between groups for each time bin.

To compare behavior performance across task learning, genotype, and actuator groups (tdTomato and ChrimsonR), we used separate two-factor mixed effects models for control and R6/2 mice, with the fixed effect terms for actuator and session number, the interaction term between actuator and session number, and a random effect term grouped by animal (Figures 4b, 4c, 4d, 4e).

$$y \sim actuator + session\ number + actuator{:}session\_number + (1\ |animal)$$

With fixed main effect terms for actuator (tdT, ChR) and session number. For Fig. 4f and Extended Fig. 5b, the fixed main effect was trial type (light, no light).
If a significant interaction was detected, posthoc pairwise Mann-Whitney U tests were performed between actuator groups for each session number.
All statistical test results are summarized in Extended Data Table 1.

### References:


33. Mangiarini, L. *et al.* Exon 1 of the HD Gene with an Expanded CAG Repeat Is Sufficient to Cause a Progressive Neurological Phenotype in Transgenic Mice. *Cell* **87**, 493–506 (1996).

34. Peters, A. J., Chen, S. X. & Komiyama, T. Emergence of reproducible spatiotemporal activity during motor learning. *Nature* **510**, 263–267 (2014).

35. Holtmaat, A. *et al.* Long-term, high-resolution imaging in the mouse neocortex through a chronic cranial window. *Nat Protoc* **4**, 1128–1144 (2009).

36. Burgold, J. *et al.* Cortical circuit alterations precede motor impairments in Huntington's disease mice. *Sci Rep-uk* **9**, 6634 (2019).

37. Warren, R. A. *et al.* A rapid whisker-based decision underlying skilled locomotion in mice. *eLife* **10**, e63596 (2021).

38. Pachitariu, M. *et al.* Suite2p: beyond 10,000 neurons with standard two-photon microscopy. *Biorxiv* 061507 (2017) doi:10.1101/061507.

39. Stringer, C., Wang, T., Michaelos, M. & Pachitariu, M. Cellpose: a generalist algorithm for cellular segmentation. *Nat. Methods* **18**, 100–106 (2021).

40. Bohnslav, J. P. *et al.* DeepEthogram, a machine learning pipeline for supervised behavior classification from raw pixels. *Elife* **10**, e63377 (2021).

41. Mathis, A. *et al.* DeepLabCut: markerless pose estimation of user-defined body parts with deep learning. *Nat Neurosci* **21**, 1281–1289 (2018).

42. Machado, A. S., Darmohray, D. M., Fayad, J., Marques, H. G. & Carey, M. R. A quantitative framework for whole-body coordination reveals specific deficits in freely walking ataxic mice. *Elife* **4**, e07892 (2015).

43. Murphy, J. I., Weaver, N. E. & Hendricks, A. E. Accessible analysis of longitudinal data with linear mixed effects models. *Dis. Model. Mech.* **15**, dmm048025 (2022).

44. Yu, Z. *et al.* Beyond t test and ANOVA: applications of mixed-effects models for more rigorous statistical analysis in neuroscience research. *Neuron* **110**, 21–35 (2022).

### Extended Data Figures

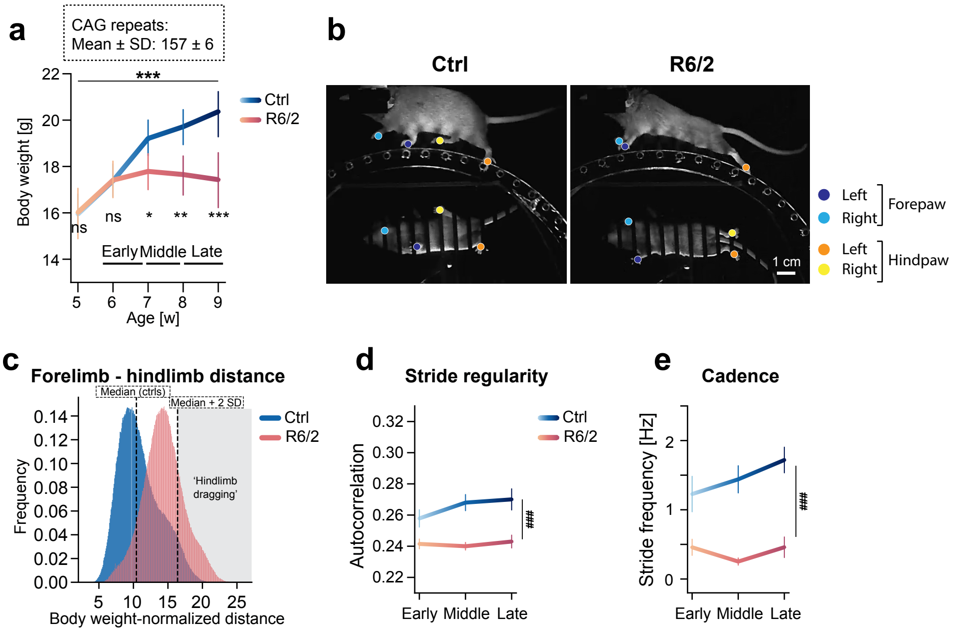


**Extended Data Fig. 1: Motor deficits in R6/2 mice during the ladder task**.
(**a**) Body weights of the R6/2 mouse colony that were used for the ladder task and had 157 ± 6 CAG repeats. Average R6/2 mouse body weight is indistinguishable from controls at the early stage of disease progression. Average weight gain slows during the middle disease stage, followed by a weight loss in the late stage in R6/2 mice. n = 45 ctrl and 61 R6/2 mice (Aligned rank transform; genotype:week interaction, $p < 0.001$, Mann-Whitney U posthoc comparisons per week: *p < 0.05, **p < 0.01, ***p < 0.001). (**b**). Example video frames of ctrl and R6/2 mice on the ladder at the middle timepoint. The trajectories of left and right fore- and hindpaws were extracted from two perspectives (side and bottom). (**c**) The weight-normalized distance between forelimbs and hindlimbs is longer in R6/2 mice due to hindlimb dragging, defined as the control median + 2 SD. (**d**) Autocorrelation of trial forelimb trajectories. The irregularity of the R6/2 gait pattern manifests as a reduction in max. autocorrelation (Aligned rank transform; genotype main effect, $p < 0.001$). (**e**) Cadence was lower in R6/2 mice (Aligned rank transform; genotype main effect, $p < 0.001$).

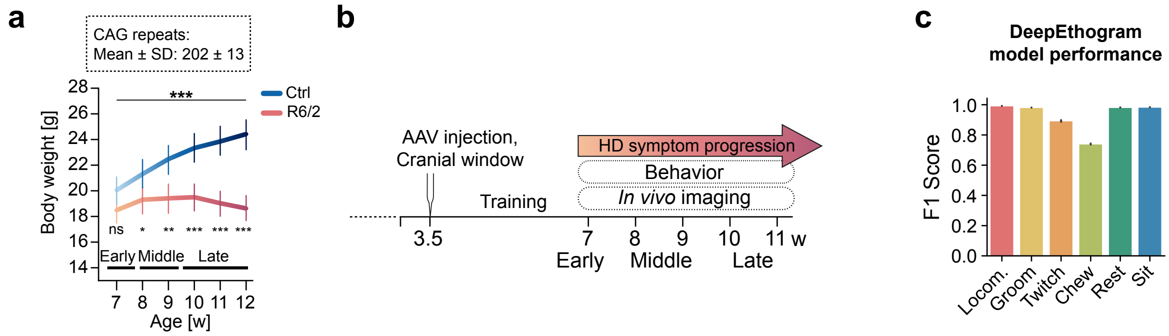


**Extended Data Fig. 2: Additional analysis of mice used for the spontaneous behavior paradigm on the wheel**.

(**a**) Body weights of the R6/2 mouse colony that were used for the wheel paradigm and had 202 ± 13 CAG repeats. ‘Early’, ‘middle’, and ‘late’ disease stages were defined as before (see Extended Data Fig. 1a; Aligned rank transform; genotype:week interaction, $p < 0.001$). (**b**) Diagram of behavioral and imaging schedule for the wheel paradigm. (**c**) Classifier model performance of DeepEthogram. F1 scores, balancing precision and recall, were calculated. Average F1 scores from 43 training iterations of the final sequence model (see Methods) are shown per behavior class.

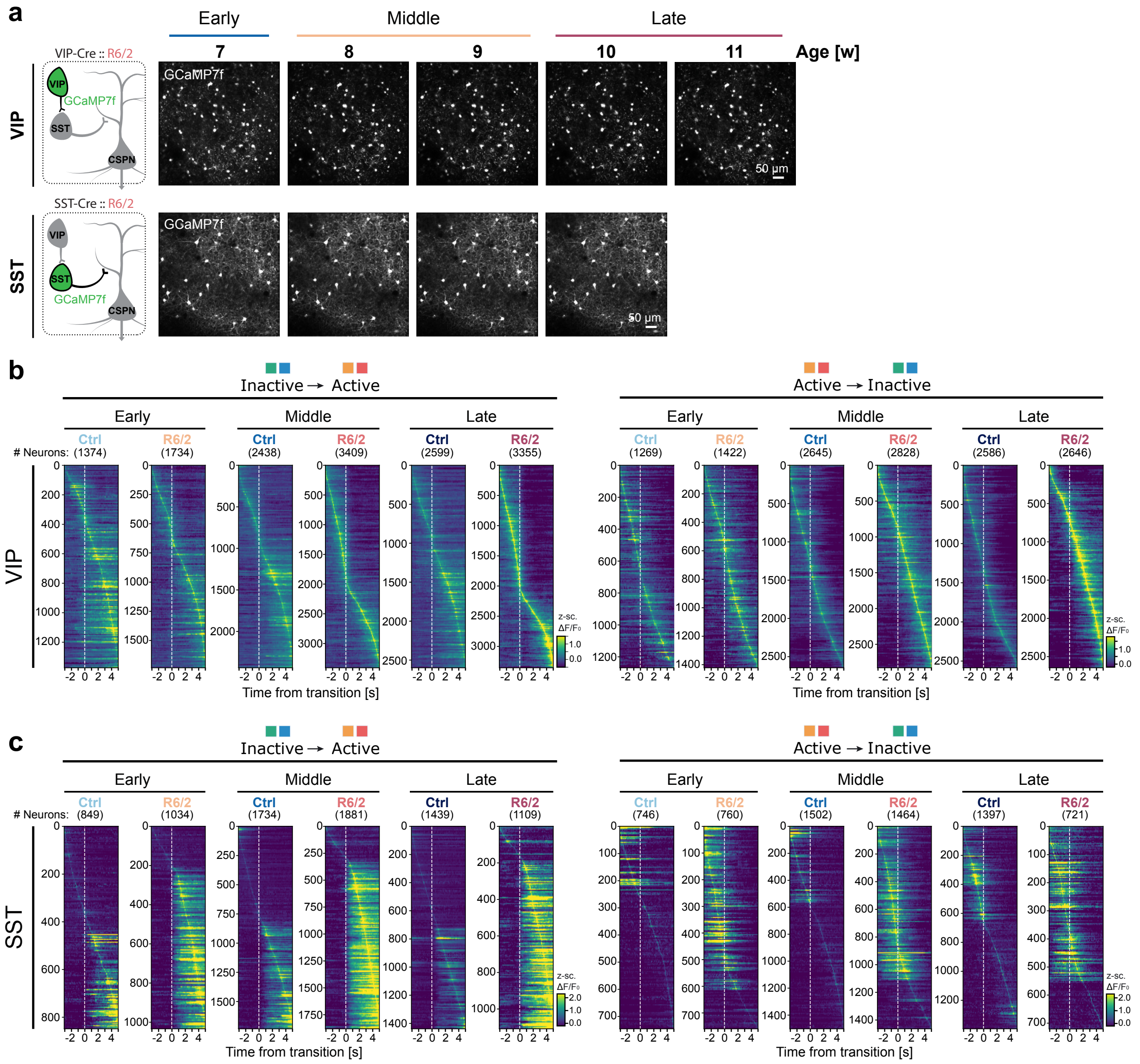


**Extended Data Fig. 3: Additional data from VIP-INs and SST-INs imaged in the wheel paradigm.**

(**a**) Example maximum intensity projections of *in vivo* two-photon fluorescence images of VIP-INs and SST-INs. Identical fields of view were imaged at weekly intervals between 7 and 10-11 weeks of age, corresponding to early, middle, and late disease stages. (**b**) z-scored $\Delta F/F_0$ from VIP-INs averaged across trials, from 3 s before behavior state transition to 5 s after (n = 8 VIP-Cre ctrl and 9 VIP-Cre::R6/2 mice). (**c**) same as (b), but for SST-INs (n = 6 SST-Cre ctrl mice and 6 SST-Cre::R6/2 mice. The numbers of neurons are indicated in the figure).

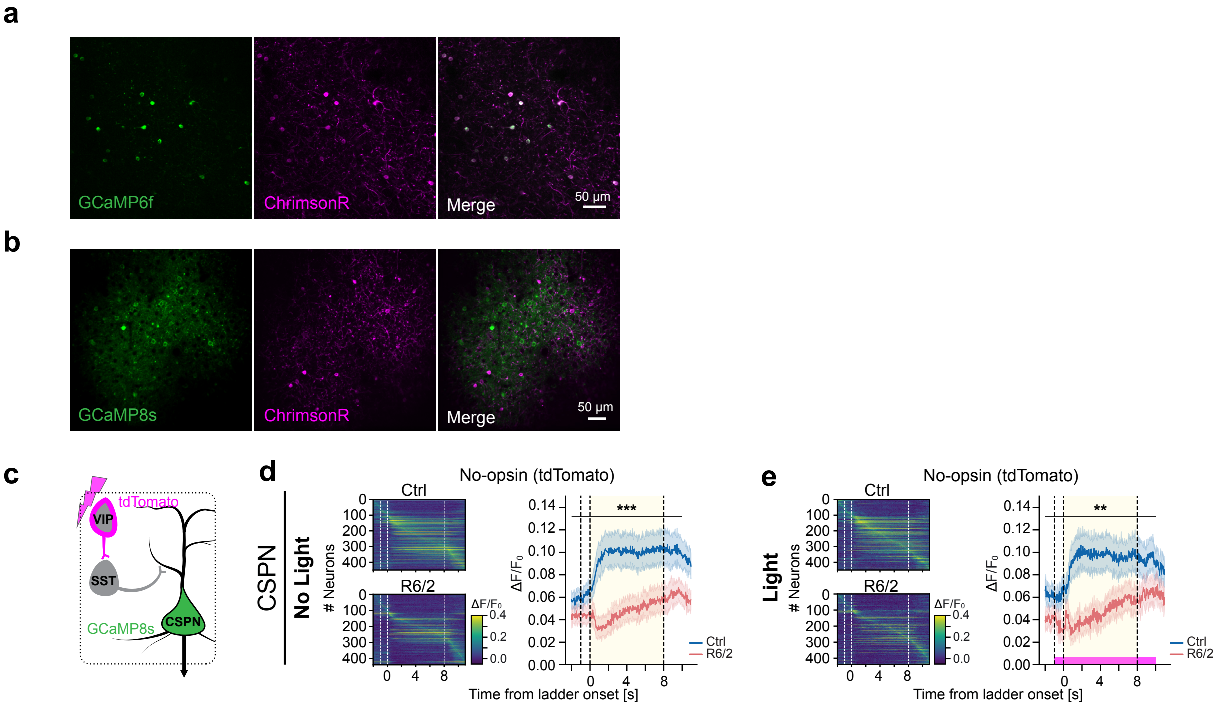


**Extended Data Fig 4: Optogenetic activation of VIP-INs normalizes VIP-IN and CSPN activity in HD mice**.

(**a-b**) Maximum intensity projections of *in vivo* two-photon fluorescence images as shown as cropped overlay images in Fig. 3a, e, detailing ChrimsonR and GCaMP6f co-expression in VIP-INs (a) and ChrimsonR expression in VIP-INs and GCaMP8s expression in CSPNs (b). (**c**) Schematic of experimental approach, expressing tdTomato in VIP-INs as no-opsin controls while imaging CSPN activity. (**d**) $\Delta F/F_0$ from CSPNs averaged across no-light trials (red masking light presented to the mouse). CSPN activity is reduced during movement in R6/2 mice. n = 450 neurons from 4 VIP-Cre ctrl mice and 440 neurons from 5 VIP-Cre::R6/2 mice (Aligned rank transform; genotype:time bin interaction, $p < 0.001$). (**e**) Same as (d), but for light trials (638 nm, 25 Hz, 2.5 mW). There is no difference in CSPN activity between light vs. no-light trials in either ctrl or R6/2 mice (Aligned rank transform; genotype:time bin interaction, $p = 0.001$).

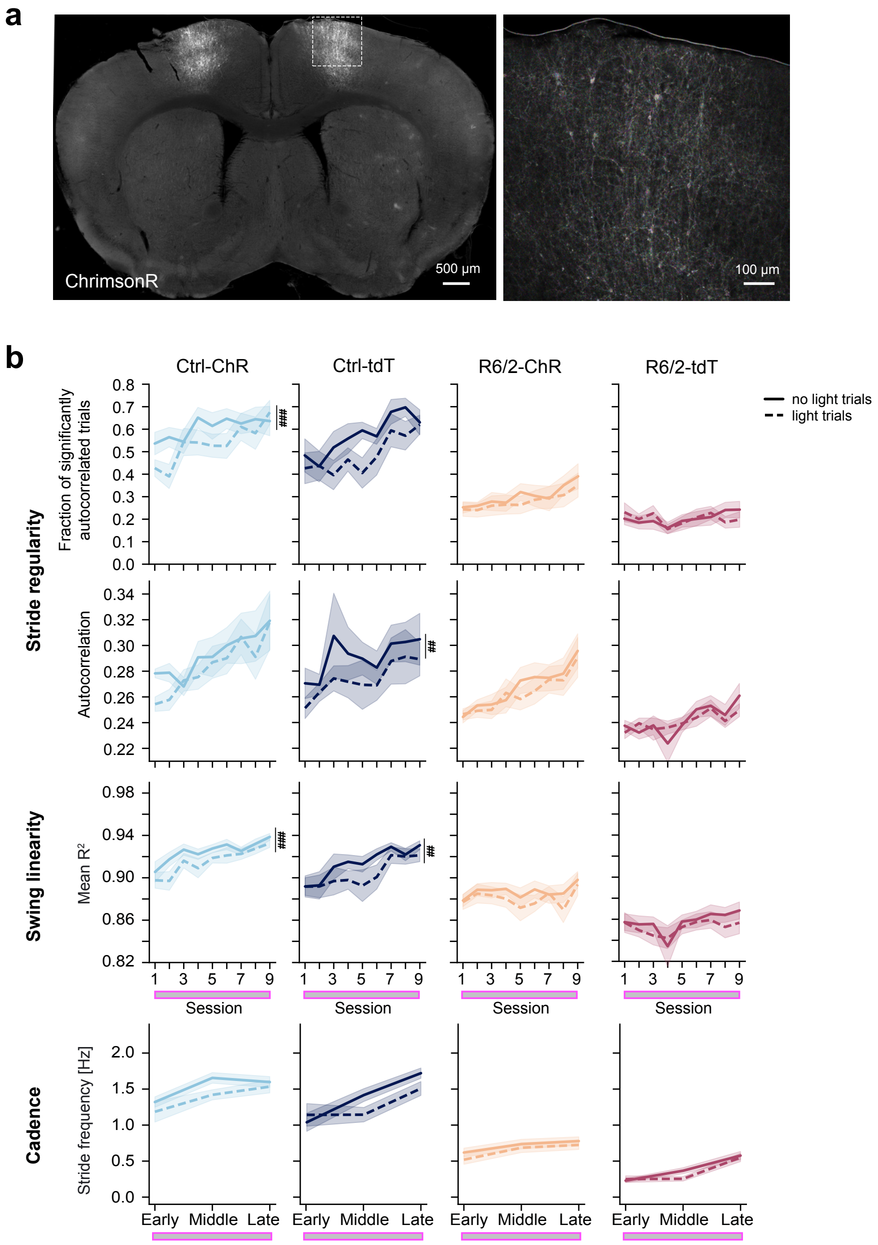


**Extended Data Fig. 5: Additional data from behavior experiments with bilateral activation of VIP-INs.**

(**a**) Example brain section from an R6/2 mouse expressing ChrimsonR in VIP-INs in the M1 cortex. Left: whole coronal section. Right: magnified image from the motor cortex, area indicated by the box on the left. (**b**) The effects of VIP-IN stimulation in R6/2 mice last beyond the light trials. Comparison of no-light and light trials for all behavioral analyses. Aligned-rank transform; Stride regularity: fraction of significantly autocorrelated trials, trial type main effect $p < 0.01$ for Ctrl-ChR, $p = 0.115$ for Ctrl-tdT, $p = 0.163$ for R6/2-ChR, $p = 1.0$ for R6/2-tdT; Autocorrelation, $p = 0.222$ for Ctrl-ChR, $p = 0.004$ for Ctrl-tdT, $p = 0.079$ for R6/2-ChR, $p = 0.513$ for R6/2-tdT. Swing linearity: trial type main effect $p < 0.001$ for Ctrl-ChR, $p = 0.004$ for Ctrl-tdT, $p = 0.052$ for R6/2-ChR, $p = 0.080$ for R6/2-tdT. Cadence: trial type main effect $p = 0.074$ for Ctrl-ChR, $p = 0.419$ for Ctrl-tdT, $p = 0.271$ for R6/2-ChR, $p = 0.206$ for R6/2-tdT. Mann-Whitney U posthoc comparisons. $p$ = n.s. for all individual sessions and all comparisons (see Extended Data Table 1). n = 9 VIP-Cre-ChR, 10 VIP-Cre-tdT, 11 VIP-Cre::R6/2-ChR, and 11 VIP-Cre::R6/2-tdT mice, 779 sessions across all mice and timepoints. Error bars represent mean ± s.e.m.

**Extended Data Table 1**

| Figure Panel | Main Factor 1 | Main Factor 1: p value | Main Factor 2 | Main Factor: 2 p value | Interaction | Interaction: p value | Posthoc pairwise comparisons: FDR-adjusted p values |
|---|---|---|---|---|---|---|---|
| Fig. 1d Dragging time / trial | genotype | 0.01168 | timepoint | 0.390722 | genotype:timepoint | 0.016909 | timepoint=early, p=0.338<br>timepoint=mid, p=0.000153<br>timepoint=late, p=7.4e-05 |
| Fig. 1e Fraction significantly autocorrelated trials | genotype | 2.4858E-06 | timepoint | 0.0043218 | genotype:timepoint | 0.0045915 | timepoint=early, p=0.000138<br>timepoint=mid, p=2.13e-08<br>timepoint=late, p=8.39e-05 |
| Fig. 1g Mean $R^2$ | genotype | 0.000029462 | timepoint | 0.657492 | genotype:timepoint | 0.036555 | timepoint=early, p=0.00273<br>timepoint=mid, p=7.11e-07<br>timepoint=late, p=3.55e-05 |
| Fig. 1i VIP | genotype | 0.018312 | time bin | 2.22E-16 | genotype:time bin | 1.7626E-12 | time bin=-2, p=0.279<br>time bin=0, p=9.36e-07<br>time bin=2, p=8.94e-10<br>time bin=4, p=7.9e-08<br>time bin=6, p=7.09e-08<br>time bin=8, p=1.16e-05 |
| Fig. 1j SST | genotype | 0.0080853 | time bin | 2.22E-16 | genotype:time bin | 2.2818E-16 | time bin=-2, p=9.75e-10<br>time bin=0, p=0.00782<br>time bin=2, p=8.39e-07<br>time bin=4, p=8.39e-07<br>time bin=6, p=8.39e-07<br>time bin=8, p=0.000138 |
| Fig. 1k CSPN | genotype | 0.00121421 | time bin | 1.9855E-06 | genotype:time bin | 0.00022688 | time bin=-2, p=0.201<br>time bin=0, p=2.28e-10<br>time bin=2, p=4.77e-08<br>time bin=4, p=4.77e-08<br>time bin=6, p=7.67e-07<br>time bin=8, p=4.72e-05 |
| Fig. 2b (Locomotion) | genotype | 1.2519E-08 | timepoint | 0.00010655 | genotype:timepoint | 2.22E-16 | timepoint=early, p=2.93e-06<br>timepoint=mid, p=2.99e-30<br>timepoint=late, p=1.43e-52 |
| Fig. 2b (Groom) | genotype | 0.0079089 | timepoint | 0.4855523 | genotype:timepoint | 0.7531268 | |
| Fig. 2b (Twitch) | genotype | 0.00004752 | timepoint | 3.1236E-10 | genotype:timepoint | 7.2683E-11 | timepoint=early, p=7.83e-05<br>timepoint=mid, p=7.56e-09<br>timepoint=late, p=2.31e-24 |
| Fig. 2b (Chew) | genotype | 0.000059926 | timepoint | 0.02373 | genotype:timepoint | 0.27093 | |
| Fig. 2b (Rest) | genotype | 0.46301 | timepoint | 1.5941E-10 | genotype:timepoint | 0.14022 | |
| Fig. 2b (Sit) | genotype | 3.5301E-10 | timepoint | 7.5213E-16 | genotype:timepoint | 2.22E-16 | timepoint=early, p=7.26e-14<br>timepoint=mid, p=5.21e-39<br>timepoint=late, p=3.36e-59 |
| Fig. 2c left panel (Early) | genotype | 0.26049 | time bin | 1.4658E-12 | genotype:time bin | 0.000062879 | time bin=-2, p=8.81e-08<br>time bin=0, p=1.03e-26<br>time bin=2, p=8.88e-21 |
| Fig. 2c left panel (Middle) | genotype | 0.073012 | time bin | 2.22E-16 | genotype:time bin | 2.22E-16 | time bin=-2, p=6.75e-19<br>time bin=0, p=8.81e-37<br>time bin=2, p=5e-71 |
| Fig. 2c left panel (Late) | genotype | 0.0096088 | time bin | 2.22E-16 | genotype:time bin | 2.22E-16 | time bin=-2, p=2.56e-46<br>time bin=0, p=3.3e-87<br>time bin=2, p=2.08e-191 |
| Fig. 2c right panel (Early) | genotype | 0.16619 | time bin | 2.22E-16 | genotype:time bin | 4.0637E-07 | time bin=-2, p=0.226<br>time bin=0, p=4.72e-13<br>time bin=2, p=4.72e-13 |
| Fig. 2c right panel (Middle) | genotype | 0.029489 | time bin | 1.0574E-15 | genotype:time bin | 2.22E-16 | time bin=-2, p=0.614<br>time bin=0, p=2.38e-21<br>time bin=2, p=3.11e-91 |
| Fig. 2c right panel (Late) | genotype | 0.50598 | time bin | 2.22E-16 | genotype:time bin | 2.22E-16 | time bin=-2, p=2.1e-59<br>time bin=0, p=0.0152<br>time bin=2, p=2.79e-50 |
| Fig. 2d left panel (Early) | genotype | 0.0050354 | time bin | 2.22E-16 | genotype:time bin | 2.22E-16 | time bin=-2, p=3.05e-07<br>time bin=0, p=6.51e-53<br>time bin=2, p=7.03e-51 |
| Fig. 2d left panel (Middle) | genotype | 0.0079475 | time bin | 2.22E-16 | genotype:time bin | 2.22E-16 | time bin=-2, p=9.04e-07<br>time bin=0, p=3.49e-113<br>time bin=2, p=1.03e-110 |
| Fig. 2d left panel (Late) | genotype | 0.041774 | time bin | 2.22E-16 | genotype:time bin | 2.22E-16 | time bin=-2, p=1.52e-08<br>time bin=0, p=3.13e-47<br>time bin=2, p=4.82e-72 |
| Fig. 2d right panel (Early) | genotype | 0.005566 | time bin | 2.22E-16 | genotype:time bin | 2.22E-16 | time bin=-2, p=7.09e-49<br>time bin=0, p=2.65e-20<br>time bin=2, p=0.211 |
| Fig. 2d right panel (Middle) | genotype | 0.03583 | time bin | 2.22E-16 | genotype:time bin | 2.22E-16 | time bin=-2, p=2.02e-85<br>time bin=0, p=3.07e-58<br>time bin=2, p=0.29 |
| Fig. 2d right panel (Late) | genotype | 0.30005 | time bin | 9.23E-11 | genotype:time bin | 2.22E-16 | time bin=-2, p=3.19e-11<br>time bin=0, p=2.87e-08<br>time bin=2, p=0.337 |
| Fig. 2e (Locomotion) | genotype | 0.020567 | timepoint | 2.22E-16 | genotype:timepoint | 3.6904E-09 | timepoint=early, p=6.02e-09<br>timepoint=mid, p=2.35e-29<br>timepoint=late, p=1.45e-82 |
| Fig. 2e (Groom) | genotype | 0.067207 | timepoint | 9.0795E-16 | genotype:timepoint | 1.0866E-11 | timepoint=early, p=6.47e-14<br>timepoint=mid, p=8.96e-38<br>timepoint=late, p=4.45e-18 |
| Fig. 2e (Twitch) | genotype | 0.10823 | timepoint | 2.22E-16 | genotype:timepoint | 1.6268E-13 | timepoint=early, p=1.16e-20<br>timepoint=mid, p=7.78e-46<br>timepoint=late, p=1.85e-13 |
| Fig. 2e (Rest) | genotype | 0.04768427 | timepoint | 2.22E-16 | genotype:timepoint | 0.00022484 | timepoint=early, p=2.18e-06<br>timepoint=mid, p=6.13e-20<br>timepoint=late, p=8.18e-34 |
| Fig. 2e (Sit) | genotype | 0.044478 | timepoint | 2.22E-16 | genotype:timepoint | 1.7542E-08 | timepoint=early, p=0.234<br>timepoint=mid, p=0.239<br>timepoint=late, p=7.16e-20 |
| Fig. 2f (Locomotion) | genotype | 0.080281 | timepoint | 2.22E-16 | genotype:timepoint | 2.22E-16 | timepoint=early, p=2.57e-58<br>timepoint=mid, p=4.05e-91<br>timepoint=late, p=5.13e-54 |

| Figure Panel | Main Factor 1 | Main Factor 1: p value | Main Factor 2 | Main Factor: 2 p value | Interaction | Interaction: p value | Posthoc pairwise comparisons: FDR-adjusted p values |
|---|---|---|---|---|---|---|---|
| Fig. 2f (Groom) | genotype | 0.12558 | timepoint | 2.0389E-15 | genotype:timepoint | 2.22E-16 | timepoint=early, p=4.06e-61<br>timepoint=mid, p=3.92e-37<br>timepoint=late, p=2.76e-20 |
| Fig. 2f (Twitch) | genotype | 0.023338 | timepoint | 7.4666E-11 | genotype:timepoint | 0.272894 | |
| Fig. 2f (Rest) | genotype | 0.782083 | timepoint | 0.016395 | genotype:timepoint | 0.412104 | |
| Fig. 2f (Sit) | genotype | 0.93257064 | timepoint | 0.00064107 | genotype:timepoint | 0.000032776 | timepoint=early, p=0.0123<br>timepoint=mid, p=0.0247<br>timepoint=late, p=0.599 |
| Fig. 3b VIP no light | genotype | 0.022802 | time bin | 2.22E-16 | genotype:time bin | 2.769E-14 | time bin=-2, p=0.152<br>time bin=0, p=2.46e-05<br>time bin=2, p=1.2e-08<br>time bin=4, p=1.14e-07<br>time bin=6, p=1.2e-08<br>time bin=8, p=6.29e-06 |
| Fig. 3c VIP light | genotype | 0.1670794 | time bin | 2.22E-16 | genotype:time bin | 0.0021307 | time bin=-2, p=0.574<br>time bin=0, p=0.000101<br>time bin=2, p=0.000157<br>time bin=4, p=0.0569<br>time bin=6, p=0.0297<br>time bin=8, p=0.0833 |
| Fig. 3d VIP no light (ctrl) vs. light (R6/2) | genotype | 0.93668 | time bin | 2.22E-16 | genotype:time bin | 0.59242 | |
| Fig. 3f CSPN no light | genotype | 0.18251 | time bin | 0.000043648 | genotype:time bin | 1.0521E-06 | time bin=-2, p=0.183<br>time bin=0, p=2.37e-12<br>time bin=2, p=1.25e-11<br>time bin=4, p=7.9e-09<br>time bin=6, p=7.21e-06<br>time bin=8, p=0.000571 |
| Fig. 3g CSPN light | genotype | 0.8099 | time bin | 0.000014358 | genotype:time bin | 0.8673 | |
| Fig. 3h CSPN no light (ctrl) vs. light (R6/2) | genotype | 0.87772 | time bin | 0.00001576 | genotype:time bin | 0.36774 | |
| Fig. 4b Dragging time / trial (ctrl) | actuator | 0.4137748 | session number | 3.5765E-14 | actuator:session number | 0.00013017 | session number=0, p=0.958<br>session number=1, p=0.414<br>session number=2, p=0.414<br>session number=3, p=0.459<br>session number=4, p=0.51<br>session number=5, p=0.51<br>session number=6, p=0.779<br>session number=7, p=0.854<br>session number=8, p=0.958<br>session number=9, p=0.854 |
| Fig. 4b Dragging time / trial (R6/2) | actuator | 0.49358628 | session number | 0.05953656 | actuator:session number | 0.00010685 | session number=0, p=0.953<br>session number=1, p=0.953<br>session number=2, p=0.953<br>session number=3, p=0.953<br>session number=4, p=0.953<br>session number=5, p=0.461<br>session number=6, p=0.461<br>session number=7, p=0.131<br>session number=8, p=0.131<br>session number=9, p=0.131 |
| Fig. 4c Fraction significantly autocorrelated trials (ctrl) | actuator | 0.5233 | session number | 3.9629E-14 | actuator:session number | 0.49225 | |
| Fig. 4c Fraction significantly autocorrelated trials (R6/2) | actuator | 0.0260291 | session number | 0.0048891 | actuator:session number | 0.0438186 | session number=0, p=0.694<br>session number=1, p=0.261<br>session number=2, p=0.0514<br>session number=3, p=0.175<br>session number=4, p=0.0209<br>session number=5, p=0.0209<br>session number=6, p=0.0725<br>session number=7, p=0.329<br>session number=8, p=0.0486<br>session number=9, p=0.0209 |
| Fig. 4c Autocorrelation (ctrl) | actuator | 0.52743 | session number | 5.2221E-07 | actuator:session number | 0.72716 | |
| Fig. 4c Autocorrelation (R6/2) | actuator | 0.0047938 | session number | 5.3185E-14 | actuator:session number | 0.0181271 | session number=0, p=0.325<br>session number=1, p=0.05<br>session number=2, p=0.00936<br>session number=3, p=0.112<br>session number=4, p=0.00936<br>session number=5, p=0.00936<br>session number=6, p=0.047<br>session number=7, p=0.05<br>session number=8, p=0.00936<br>session number=9, p=0.00936 |
| Fig. 4d Stride frequency (ctrl) | actuator | 0.4710383 | timepoint | 3.8247E-12 | actuator:timepoint | 0.0015405 | timepoint=baseline, p=0.591<br>timepoint=early, p=0.0172<br>timepoint=mid, p=0.0803<br>timepoint=late, p=0.446 |
| Fig. 4d Stride frequency (R6/2) | actuator | 0.028413 | timepoint | 7.5812E-10 | actuator:timepoint | 0.032157 | timepoint=baseline, p=0.0179<br>timepoint=early, p=6.56e-10<br>timepoint=mid, p=0.000156<br>timepoint=late, p=0.00346 |
| Fig. 4e Mean $R^2$ (ctrl) | actuator | 0.25225 | session number | 2.22E-16 | actuator:session number | 0.14599 | |
| Fig. 4e Mean $R^2$ (R6/2) | actuator | 0.0019844 | session number | 0.0031532 | actuator:session number | 0.0195846 | session number=0, p=0.212<br>session number=1, p=0.0177<br>session number=2, p=0.000874<br>session number=3, p=8.7e-05<br>session number=4, p=0.000512<br>session number=5, p=0.00151<br>session number=6, p=0.00724<br>session number=7, p=0.00831<br>session number=8, p=0.0278<br>session number=9, p=0.00134 |

| Figure Panel | Main Factor 1 | Main Factor 1: p value | Main Factor 2 | Main Factor: 2 p value | Interaction | Interaction: p value | Posthoc pairwise comparisons: FDR-adjusted p values |
|---|---|---|---|---|---|---|---|
| Fig. 4f Dragging time / trial (ctrl-ChR) | trial type | 1.8278E-07 | session number | 0.262873 | trial type:session number | 0.015817 | session number=1, p=0.926<br>session number=2, p=0.926<br>session number=3, p=0.926<br>session number=4, p=0.926<br>session number=5, p=0.926<br>session number=6, p=0.926<br>session number=7, p=0.926<br>session number=8, p=0.926<br>session number=9, p=0.926 |
| Fig. 4f Dragging time / trial (ctrl-tdT) | trial type | 0.51008 | session number | 1.6632E-06 | trial type:session number | 0.80906 | |
| Fig. 4f Dragging time / trial (R6/2-ChR) | trial type | 0.7264369 | session number | 0.00071113 | trial type:session number | 0.99896398 | |
| Fig. 4f Dragging time / trial (R6/2-tdT) | trial type | 0.885325 | session number | 0.090863 | trial type:session number | 0.999374 | |
| Fig. S1a Body weight | genotype | 0.015049 | week | 2.22E-16 | genotype:week | 2.22E-16 | week=5.0, p=0.968<br>week=6.0, p=0.968<br>week=7.0, p=0.0299<br>week=8.0, p=0.00141<br>week=9.0, p=0.000955 |
| Fig. S1d Autocorrelation | genotype | 0.00028384 | timepoint | 0.09291093 | genotype:timepoint | 0.1137976 | |
| Fig. S1e Stride frequency | genotype | 0.00016598 | timepoint | 0.01935397 | genotype:timepoint | 0.07510544 | |
| Fig. S2a Body weight | genotype | 0.000044848 | week | 2.22E-16 | genotype:week | 2.22E-16 | week=7, p=0.061<br>week=8, p=0.0286<br>week=9, p=0.00259<br>week=10, p=0.000216<br>week=11, p=6e-06<br>week=12, p=2e-06 |
| Fig. S4d CSPN no light (no-opsin) | genotype | 0.024623 | time bin | 0.000024299 | genotype:time bin | 4.8752E-06 | time bin=-2, p=0.00058<br>time bin=0, p=4.41e-17<br>time bin=2, p=2.89e-15<br>time bin=4, p=1.74e-11<br>time bin=6, p=2.45e-09<br>time bin=8, p=1.25e-05 |
| Fig. S4e CSPN light (no-opsin) | genotype | 0.02907875 | time bin | 0.00020829 | genotype:time bin | 0.00103343 | time bin=-2, p=4.98e-06<br>time bin=0, p=1.55e-15<br>time bin=2, p=1.15e-13<br>time bin=4, p=2.41e-11<br>time bin=6, p=2.82e-07<br>time bin=8, p=0.000225 |
| Fig. S5b Fraction significantly autocorrelated trials (ctrl_ChR) | trial type | 0.000086124 | session number | 0.000023466 | trial type:session number | 0.065401 | session number=1, p=0.382<br>session number=2, p=0.339<br>session number=3, p=0.94<br>session number=4, p=0.382<br>session number=5, p=0.382<br>session number=6, p=0.382<br>session number=7, p=0.668<br>session number=8, p=0.7<br>session number=9, p=0.517 |
| Fig. S5b Fraction significantly autocorrelated trials (ctrl_tdT) | trial type | 0.00019861 | session number | 2.6484E-06 | trial type:session number | 0.56715807 | session number=1, p=0.766<br>session number=2, p=0.965<br>session number=3, p=0.565<br>session number=4, p=0.766<br>session number=5, p=0.195<br>session number=6, p=0.565<br>session number=7, p=0.766<br>session number=8, p=0.565<br>session number=9, p=0.965 |
| Fig. S5b Fraction significantly autocorrelated trials (R6/2_ChR) | trial type | 0.16351 | session number | 0.03149 | trial type:session number | 0.98337 | |
| Fig. S5b Fraction significantly autocorrelated trials (R6/2_tdT) | trial type | 0.99988 | session number | 0.2409 | trial type:session number | 0.63321 | |
| Fig. S5b Autocorrelation (ctrl_ChR) | trial type | 0.2221922 | session number | 0.0002171 | trial type:session number | 0.3773171 | |
| Fig. S5b Autocorrelation (ctrl_tdT) | trial type | 0.0040582 | session number | 0.0287818 | trial type:session number | 0.9820537 | session number=1, p=0.604<br>session number=2, p=0.604<br>session number=3, p=0.604<br>session number=4, p=0.604<br>session number=5, p=0.604<br>session number=6, p=0.604<br>session number=7, p=0.604<br>session number=8, p=0.604<br>session number=9, p=0.724 |
| Fig. S5b Autocorrelation (R6/2_ChR) | trial type | 0.079101 | session number | 2.3466E-07 | trial type:session number | 0.895923 | |
| Fig. S5b Autocorrelation (R6/2_tdT) | trial type | 0.51293 | session number | 0.000043263 | trial type:session number | 0.78056 | |
| Fig. S5b Mean $R^2$ (ctrl_ChR) | trial type | 0.00014138 | session number | 2.0821E-10 | trial type:session number | 0.77426355 | session number=1, p=0.669<br>session number=2, p=0.669<br>session number=3, p=0.669<br>session number=4, p=0.669<br>session number=5, p=0.669<br>session number=6, p=0.669<br>session number=7, p=0.91<br>session number=8, p=0.762<br>session number=9, p=0.669 |
| Fig. S5b Mean $R^2$ (ctrl_tdT) | trial type | 0.0039846 | session number | 1.0874E-08 | trial type:session number | 0.8373335 | session number=1, p=0.93<br>session number=2, p=0.93<br>session number=3, p=0.93<br>session number=4, p=0.71<br>session number=5, p=0.93<br>session number=6, p=0.71<br>session number=7, p=0.93<br>session number=8, p=0.93<br>session number=9, p=0.868 |
| Fig. S5b Mean $R^2$ (R6/2_ChR) | trial type | 0.0525136 | session number | 0.0085335 | trial type:session number | 0.8075818 | |
| Fig. S5b Mean $R^2$ (R6/2_tdT) | trial type | 0.079881 | session number | 0.028338 | trial type:session number | 0.973319 | |
| Fig. S5b Stride frequency (ctrl_ChR) | trial type | 0.07417538 | timepoint | 0.00018849 | trial type:timepoint | 0.4242176 | |
| Fig. S5b Stride frequency (ctrl_tdT) | trial type | 0.41892 | timepoint | 9.4304E-09 | trial type:timepoint | 0.17018 | |
| Fig. S5b Stride frequency (R6/2_ChR) | trial type | 0.2709024 | timepoint | 0.0041379 | trial type:timepoint | 0.8453192 | |
| Fig. S5b Stride frequency (R6/2_tdT) | trial type | 0.20625 | timepoint | 5.776E-11 | trial type:timepoint | 0.32078 | |
